# Expansion load reduces fitness at the range margin of an invasive plant

**DOI:** 10.64898/2026.07.31.741867

**Authors:** Ryan Briscoe Runquist, John W. Benning, David A. Moeller

## Abstract

Genetic drift and natural selection jointly shape how populations evolve during range expansion, yet their joint influence on fitness at expanding range margins is rarely examined. Serial founder events during range expansion may amplify genetic drift and result in the accumulation (or greater expression) of deleterious mutations at range margins, termed expansion load. Conversely, natural selection should lead to local adaptation during expansion provided that populations harbor adequate genetic variation. We used a manipulative transplant experiment, genome-wide SNP data, and demographic surveys of the invasive plant, common tansy (*Tanacetum vulgare*), to evaluate evidence for expansion load and local adaptation during ca. 150 years of invasion in Minnesota, U.S.A. The common garden occurred near the southern range margin and included 16 populations that span (1) a chronosequence of invasion from northeastern MN (invasion core) to southern and western range margins and (2) environmental gradients in temperature and precipitation. We also manipulated temperature and precipitation to test whether environmental stress amplified the expression of expansion load or revealed population differentiation in climate adaptation. Population fitness declined strongly with distance from the invasion core, consistent with the accumulation of expansion load. This finding was corroborated by analyses of 173 populations that showed an increase in homozygosity (*F*) and a decrease in population size from the invasion core to range margins. We did not find any evidence of local adaptation; a strong positive relationship between population mean fitness and environmental distance was indicative of maladaptation. While the elevated temperature manipulation reduced fitness, it did not amplify the expression of expansion load. Taken together, our results are consistent with the hypothesis that serial population bottlenecks and strong genetic drift led to expansion load and reduced fitness at range margins.

**Teaser Text:** Species’ geographic ranges are dynamic over geologic time scales but may also shift rapidly in response to climate change and for invasive species. While range shifts are often viewed only through an ecological lens, the extent and pace of range expansion may be significantly modulated by evolutionary processes. We disentangled the influence of genetic drift and natural selection on population fitness across an invasion chronosequence (spanning ca. 150 years) for the herbaceous plant common tansy (*Tanacetum vulgare*). Our synthesis of analyses of a field transplant experiment, genome-wide SNP data, and demographic surveys provided evidence consistent with the accumulation of expansion load at leading range edges. While theory often predicts that local adaptation should occur readily in response to novel environments, our results were consistent with local maladaptation. Our findings emphasize the importance of stochastic processes in shaping the geographic distribution of species and their capacity to shift with changing environments.

## Introduction

Contemporary range expansions are valuable opportunities to investigate how populations evolve over short time periods in response to spatial and temporal environmental variation (Colautti & Lau, 2015; Sexton et al., 2009). Evolutionary theory has shown that the interplay of genetic drift and natural selection is pivotal in modulating range expansion (Angert et al., 2020; Miller et al., 2020). Serial founder events during range expansion may amplify genetic drift thereby limiting the capacity for local adaptation (Polechová, 2018) and promoting the accumulation of genetic load (Peischl et al., 2015). However, experimental studies have rarely simultaneously tested for evidence of local adaptation and accumulating genetic load in the process of range expansion in wild systems.

During range expansion, repeated founding events by small groups of individuals causes genetic drift to be a powerful force and results in low effective population sizes and reduced efficacy of selection at the advancing front (Peischl et al., 2013; Peischl & Excoffier, 2016). While drift often causes losses of genetic variation, it may also cause some alleles to rise to high frequency purely due to stochastic sampling. In the context of range expansion, this process is often termed allele surfing (Excoffier & Ray, 2008; Klopfstein et al., 2006). When these surfing alleles are deleterious, the reduced efficacy of purifying selection at the advancing front allows them to accumulate, a phenomenon often called expansion load. Expansion load may depress fitness at range margins and can persist well behind the advancing front (Gilbert et al., 2018; Peischl et al., 2013, 2015; Perrier et al., 2020). Expansion load accrues fastest where newly established populations grow rapidly from low density, which reduces the efficacy of selection (Klopfstein et al., 2006; Peischl et al., 2015). Additionally, small, mate-limited founding populations may experience inbreeding during establishment, and any partially recessive deleterious alleles masked in heterozygotes in the core may become expressed in homozygotes at the front, further increasing genetic load (Peischl & Excoffier, 2015; Schrieber & Lachmuth, 2017; Uller & Leimu, 2011).

Direct evidence that expansion load depresses fitness in wild populations remains limited. Genomic signatures consistent with accumulating deleterious mutations have been detected during range expansions in both humans (Henn et al., 2016; Peischl et al., 2018) and plants (González-Martínez et al., 2017). Laboratory experiments have shown that expansion generates load under controlled conditions (Bosshard et al., 2017; Holtz et al., 2025; Weiss-Lehman et al., 2017). The few studies directly documenting a fitness cost in wild populations have focused on species that expanded from Pleistocene glacial refugia, producing strong evidence of accumulated load across this historical expansion (Koski et al., 2019; Perrier et al., 2020). But whether expansion load mediates fitness across a contemporary, actively expanding range remains untested.

As a species establishes across heterogeneous environments, natural selection should lead to local adaptation if populations harbor adequate genetic variation for fitness (Holt, 2003; Kirkpatrick & Barton, 1997; Miller et al., 2020). Field transplant experiments provide insight into the degree of local adaptation. One approach tests whether fitness declines with environmental divergence between the geographic origin of the genotype and the experimental site, consistent with local adaptation (Gorton et al., 2022; Peschel et al., 2025). If this relationship is absent or inverted (i.e. maladaptation), several scenarios could be at play: 1) the invasion is relatively recent and there has been insufficient time for local adaptation to have occurred, 2) strong gene flow from core populations has constrained the response to selection in edge populations, 3) or selection is weak and/or inefficient relative to the strength of genetic drift.

Environmental variation across the landscape often mediates fitness during range expansion. Even in the presence of expansion load, populations may locally adapt as the expansion front enters novel environments (Peischl et al., 2015). At the same time, stressful environments at range margins may amplify expansion load (Perrier et al., 2022). Manipulating environments in a common garden is a particularly powerful approach to testing whether specific environmental factors drive local adaptation or modulate the expression of expansion load. For example, if populations have locally adapted to climatic gradients, they should respond to climatic treatments differently based on their source environment. Local adaptation to the manipulated variable is expected to manifest as relative increases in fitness for populations brought nearer to their home-site conditions by the manipulation, and relative decreases in fitness for populations facing increased mismatch due to the manipulation. If populations are subject to expansion load, such environmental stress may amplify fitness losses for range edge populations carrying elevated load — a phenomenon we refer to as stress-amplified expansion load (sensu Cheptou et al., 2000; Fox & Reed, 2011; Perrier et al., 2022). Conducting environmental manipulations with populations sourced from across the range can thus sharpen predictions about population responses to environmental change, and how they may vary with range position (Gilbert et al., 2017, 2018).

Experiments with invasive species can help disentangle the relative influence of natural selection and genetic drift on evolution during range expansion (Clements & Jones, 2021; Colautti & Lau, 2015). While studies of post-glacial migrations illuminate the long-term consequences of range expansions (Hewitt, 2000; Perrier et al., 2020), post-expansion gene flow, selection, and drift may obscure how evolutionary dynamics influenced populations during the active expansion period (Excoffier et al., 2009). In many invasive species, precise information is available on the timing of introduction and the location of range cores and actively advancing range edges (Colautti & Lau, 2015), which facilitates tests for expansion load. Adaptive evolution in invasive species was originally thought to be constrained by reduced genetic variation caused by frequent population bottlenecks (Baker & Stebbins, 1965; Hodgins et al., 2025). However, multiple introductions and post-introduction admixture have resulted in substantial genetic variation in many invasive species (Dlugosch & Parker, 2008; Estoup et al., 2016; Schrieber & Lachmuth, 2017). Consequently, adaptive evolution has been widely documented in invasive species populations (Hodgins et al., 2025; Oduor et al., 2016; van Kleunen et al., 2018), including the rapid recapitulation of clinal patterns in the native range (Colautti et al., 2009; Colautti & Lau, 2015). This focus on adaptation has, however, left expansion load largely unexamined in contemporary invasions.

We tested for evidence of expansion load and local adaptation in the range expansion of common tansy (*Tanacetum vulgare*) in Minnesota, U.S.A (MN). Common tansy is an invasive plant of the northern United States that has expanded its range dramatically within the last ca. 150 years. This range expansion has occurred along pronounced gradients in both temperature and precipitation from cooler, mesic northeastern habitats into warmer, drier regions at the southern and western range edges. We conducted a field transplant experiment over two years that included 16 source populations that span the chronosequence of range expansion and environmental gradients in MN. We found substantial population differentiation in mean fitness, suggesting that evolution has occurred rapidly during range expansion. We then examined the extent to which population fitness variation was explained by distance to the range core (expansion load) versus distance in environment between source populations and the transplant site [local (mal)adaptation]. Our analyses revealed strong evidence for a decline in fitness from the core to margins of the invasion chronosequence. We further tested the expansion load hypothesis by asking whether population sizes declined and genome-wide homozygosity (*F*) increased with range expansion. Last, we used experimental manipulations of temperature and precipitation to ask whether environmental stress amplified expansion load and whether there was evidence for local adaptation to these specific climatic gradients.

## Methods

### Natural history

#### Species background

Common tansy (*Tanacetum vulgare* L.; Asteraceae) is an herbaceous, short-lived perennial native to Eurasia (Fig. 1). Individuals are largely self-incompatible and outcrossing is facilitated by diverse insect pollinators (LeCain & Sheley, 2014; Lokki et al., 1973); however, self-fertilization may occur rarely (Frank & Klotz, 1988; Prach & Wade, 1992). Recruitment occurs through seed, rhizomes, and root fragments, with dispersal occurring via wind, water, and anthropogenic activity (LeCain & Sheley, 2014; White, 1997). Because it produces a diverse set of toxic terpenes, *T. vulgare* is avoided by livestock and most insect herbivores, enhancing its capacity to outcompete neighboring plants in grasslands and pastures (Wolf et al., 2012).

**Figure 1.**
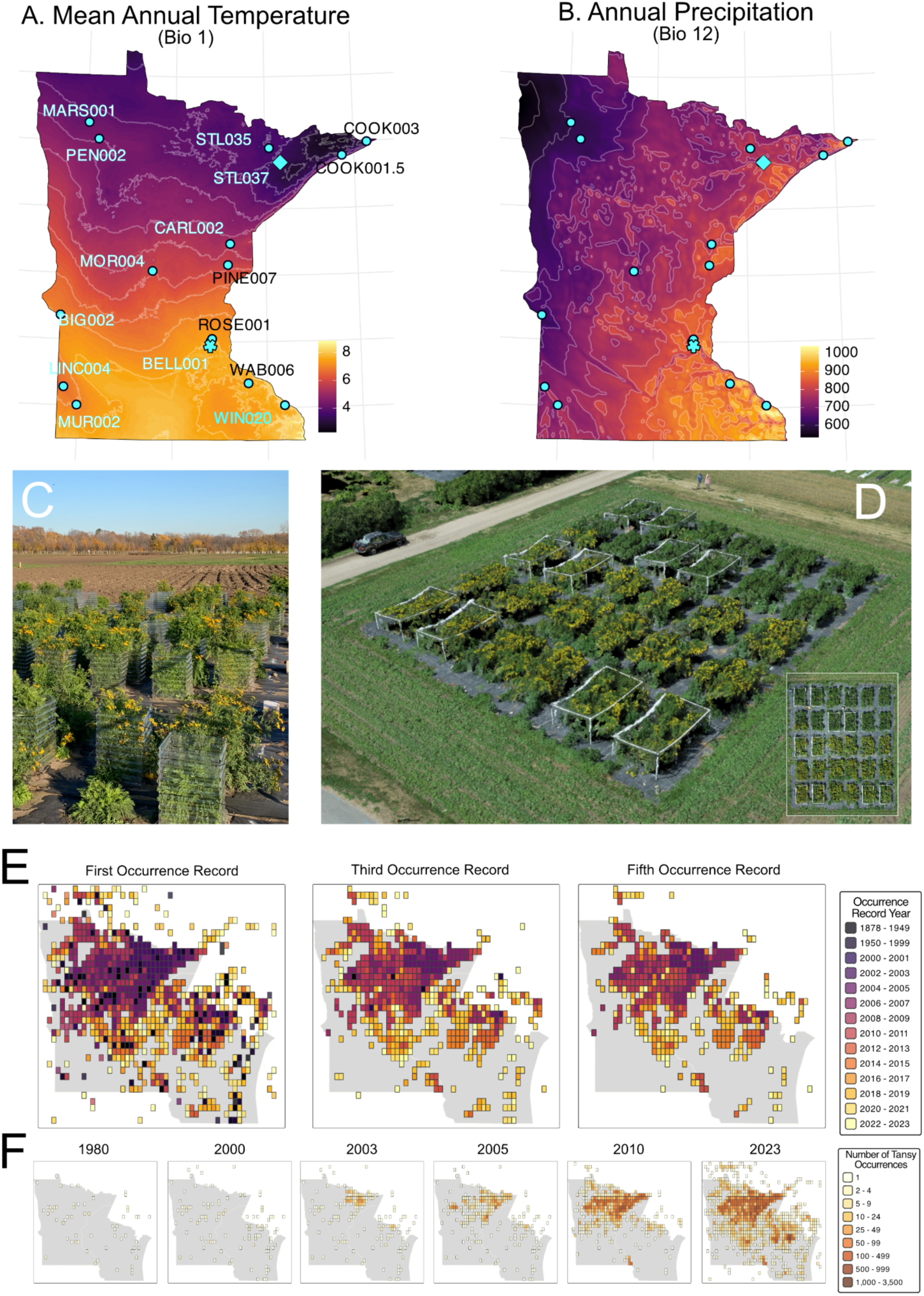
Broad scale climate gradients across Minnesota: A) Mean Annual Temperature (Bio 1) and B) Annual Precipitation (Bio 12). A & B) Common garden source population localities (cyan circles), experimental garden site (cyan star) and the locality of the invasion core (cyan diamond). C) Image of common garden during harvest 2021 with heat shelters (OTCs) in place. D) Aerial image of experimental common garden taken from drone imaging; inset is birds-eye view of entire field plot. E & F) Chronological spread of common tansy in MN: E) Gridded year of first, third, and fifth occurrence record - GBIF + EDDMaps. Darker colors indicate earlier record date; F) Accumulation of common tansy occurrence records 1980 – 2023. Darker grid cells indicate more occurrence records. **Alt. Text**: Maps and pictures with subpanels A-F showing the major temperature and precipitation environmental gradients with points at the map positions of the source populations and experimental site, pictures of the experimental design, and a time-series of common tansy invasion across Minnesota.

#### Invasion history and climate variation

Introduced to the northeastern U.S. in the 1600s, common tansy is now distributed across the northern United States, with the highest recorded densities occurring in Minnesota, U.S.A. (Briscoe Runquist & Moeller, 2024; Lake et al., 2020; Mack, 2003; Mitich, 1992; Roberts, 1878). In Minnesota (MN), the timing and accumulation of occurrence records indicates that common tansy spread rapidly from its core invaded area in northeastern MN (since ca. 1875; (Mack, 2003)) to southern and western regions, particularly over the last two decades (Fig. 1E & F) (Briscoe Runquist & Moeller, 2024; Jacobs, 2008; LeCain & Sheley, 2014) (Fig. 1). The regional climate is characterized by a latitudinal temperature gradient and a longitudinal precipitation gradient (declining from east to west). Under future climate scenarios, the region is projected to experience increased mean temperatures and greater unpredictability in spring and summer precipitation events: increased mean precipitation but longer periods of drought during the growing season (Chen & Ford, 2023; Ford et al., 2021; Kunkel et al., 2022; Liess et al., 2022)(Fig. 1A & B).

### Population seed collection

In fall 2019, we collected seeds from 16 populations that span regional temperature and precipitation gradients (Fig. 1A). These populations also span areas that include recent and rapid range expansion (Fig. 1). Notably, population STL037 lies closest to the core invaded area where *T. vulgare* first became highly abundant. Similarly, STL035, COOK001.5, and COOK003 were collected within the same northeastern invasion core region. We also collected populations that span from core to marginal areas where recent range expansion has been observed: southwest (MUR002, BIG002, LINC004), northwest (MARS001, PEN002), and southeast (WAB006, WIN020) (Fig. 1). Source populations along this chronosequence of invasion allowed us to test the hypothesis that expansion load influences fitness at range margins. Our collections also occurred in a continuous fashion with respect to environmental distance from the transplant site. For example, BELL001 is a natural population very close to the experimental site and would be expected to have highest fitness under a hypothesis of local adaptation. Importantly, our population sampling and placement of the transplant site resulted in weak correlations between environmental distance (from transplant site to source population) and distance to core (r = −0.5, *P* = 0.051).

In each source population, we collected seeds from ca. 50 inflorescences from each of ca. 10 maternal plants (mean: 7.5) spaced at least 10 m apart to minimize sampling clones or close relatives. Variation in the number of maternal families collected reflects variation in population size. We pooled seeds from each population, including an equal mass of seeds from maternal families.

### Population demographic and genetic characteristics

Source populations for this experiment are a subset of a broader statewide survey of 173 populations (Table S1). At each survey site, we estimated population size using an ordinal categorical scale (1 = 1-10 individuals, 2 = 10-100, 3 = 100-1000, 4 = 1000-10000, 5 = 10000+). We also collected leaf tissue and stored samples in silica desiccant. One individual per population was sequenced using reduced representation sequencing, which generated a SNP dataset including 3071 loci (Briscoe Runquist & Moeller, 2024). We estimated the fixation index (***F***; also referred to as inbreeding coefficient) for each population using Plink (Purcell, 2019; Purcell et al., 2007). ***F*** quantifies excess homozygosity compared to Hardy-Weinberg expectation for a panmictic population (*F* = (H_exp_ - H_obs_)/H_exp_); we did not define a reference population for this analysis so all estimates are relative values (Yang et al., 2011). We included loci that were present in 70% of all individuals and with a minor allele frequency of ≧0.01. Individuals from BELL001, COOK001.5, and ROSE001 were not available when sequencing occurred.

### Field experiment

We conducted a transplant experiment at the Minnesota Agricultural Experiment Station near the southern range margin in MN (St. Paul, MN; Fig. 1) from spring 2021 through late fall 2022. We planted 450 seedlings into 15 blocks, each subdivided into two equal plots of 15 plants (Fig. S1). We assigned one plant per population to each plot (2 plants/population/block), with the exception of COOK001.5 and COOK003, which were represented by one plant per block. We mitigated invasion risk by planting into sunken nursery pots and surrounding experimental pots with landscape fabric (See Supplementary Material).

### Climate manipulations

We manipulated temperature and precipitation using a split-plot design. We increased temperature using open-topped containers (OTCs) constructed from clear corrugated plastic (1.5’ wide x 1.5’ deep x 2’ tall; Fig. 1). OTCs were deployed around individual plants three weeks after transplantation. In each block, one plant per population was randomly assigned to the elevated temperature treatment while the other experienced ambient temperatures. OTCs passively raised temperatures around the plant by 1-3 °C during the day and 1-2 °C during the night (Fig. S2).

We applied three precipitation treatments at the block level: ambient, reduced, and increased precipitation (5 blocks/precipitation treatment). To reduce precipitation (potentially simulating drought), we excluded ca. 30% of growing-season precipitation using event-based rain-out shelters (Fig. S3). To increase precipitation, we added water to pots within 24 hours (typically <12 h) after each rain event (See Supplementary Material). We estimated the event precipitation using measurements from the UMN climate observatory (*MN* Dept. of Natural Resources, 2021-2022), which was within 300 meters of the site, and calculated the volume of water needed to increase precipitation by 30%.

### Fitness

We assessed fitness by separately harvesting all reproductive (flowers/fruits) and vegetative aboveground biomass at the end of each growing season and dried them at 35°C prior to weighing. To account for the correlation between reproductive and vegetative biomass (r = 0.51, P < 0.001; Fig. S4), we generated a composite fitness score to estimate lifetime fitness using a Principal Component Analysis that combined cumulative reproductive and vegetative biomass for two growing seasons (See Supplementary Material). PC1 explained 75% of the variation in combined biomass and was used as our fitness metric after adding a constant to avoid negative values.

### Calculation of distance metrics

We calculated two distance metrics for each source population in the experiment: geographic distance from the invasion core and environmental distance from the common garden. We calculated geographic distances from the invasion core using the great-circle distance (via the Haversine equation) between each sampled population and the STL037 population, which is closest to the invasion core. Similarly, we calculated the environmental distance between each population source site and the population closest to the transplant site: BELL001 (Fig. 1A). We used thirteen variables from the CHELSA-bioclim and climatology datasets (Brun et al., 9/2025) annual averages and ranges (temperature, precipitation, climate moisture index), potential evapotranspiration (mean and range), frost change frequency, growing degree days (10°C), and growing-season specific attributes (first day, length, precipitation, temperature) (Table S2; Fig. S5). All climate variables were centered and scaled across the 16 populations. Environmental distance was the Euclidean distance in this z-scored space between each population and the home-site population (BELL001). We calculated three environmental distances for each population: 1) overall environmental distance (all variables), 2) temperature distance, and 3) precipitation distance (See Table S2 for details). We estimated correlations among distance metrics using Pearson correlation coefficients.

Distance from the invasion core was negatively correlated with temperature distance (r = −0.70, *P* = 0.003) but not with precipitation distance (r = 0.02, *P* = 0.955). Overall environmental distance was positively correlated with both temperature (r = 0.95, *P* < 0.0001) and precipitation distance (r = 0.82, *P* < 0.0001). Temperature and precipitation distances were also positively correlated (r = 0.61, *P* = 0.013) (Table S3).

### Statistical analyses

All analyses were conducted in R version 4.5.1 including the following packages: tidyverse for data manipulation (Wickham et al., 2019), ggplot2 for figures (Wickham, 2016), the maps package to display geographical maps (Becker et al., 2025), and geosphere for calculating geographic distances (Hijmans, 2026). Statistical analyses were conducted in mgcv (Wood, 2017) using a single modeling framework (described below). Distance variables were centered and scaled to unit variance before model fitting. We evaluated all tests at α = 0.05. We used Anthropic’s Claude Opus 4.8 and Google’s Gemini 3.6 Flash LLM’s to build and check functions, unit test, format code, improve documentation, and suggest text edits for clarity. All code and text was independently verified by the authors.

### Modeling framework

To determine the relative influence of local adaptation and expansion load on fitness within a single inferential framework, we fit fourteen candidate additive mixed models (GAMs) in mgcv. We used additive models because they allowed us to incorporate individual estimates of fitness and potential non-linear relationships between fitness and predictors. GAMs capture linear trends with parametric terms while simultaneously using flexible smooths (i.e. splines) to model non-linearity. All models were estimated with maximum likelihood (method = “ML”) because AIC values derived from ML likelihoods are directly comparable across models that differ in which fixed effects are included/excluded. Every candidate model shared random intercepts for population, population × temperature treatment and population × precipitation treatment interactions (the omnibus G × E variance components), and block (the whole-plot error term for the split-plot design) (Table 1).

**Table 1.**
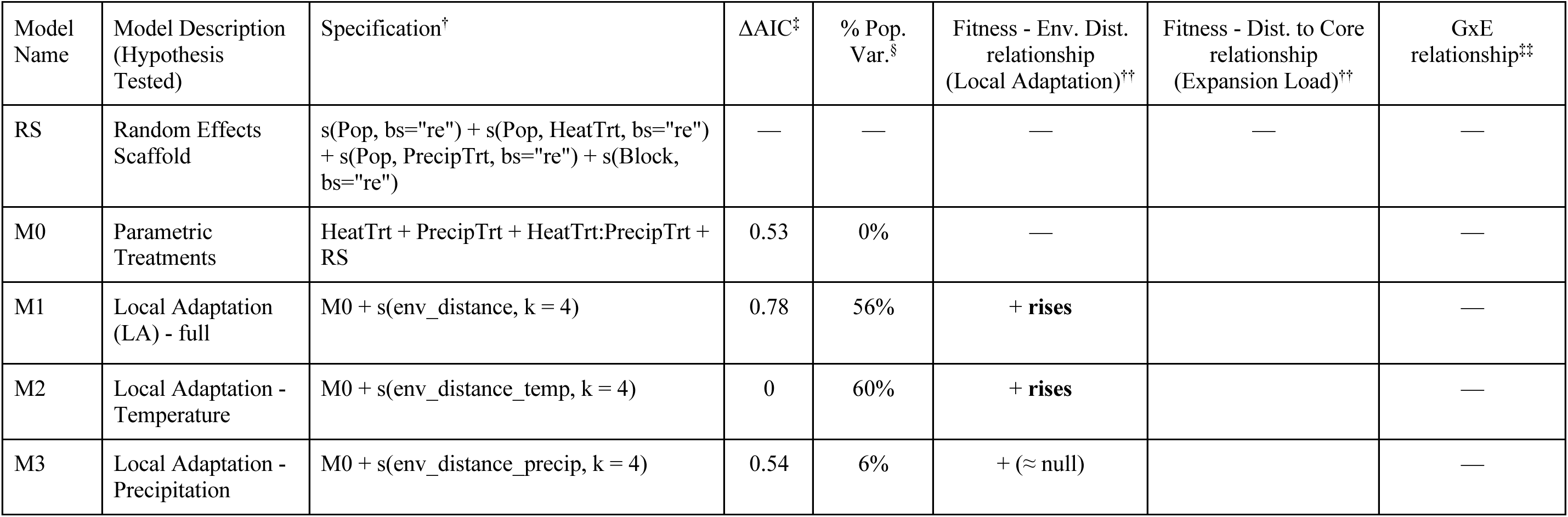
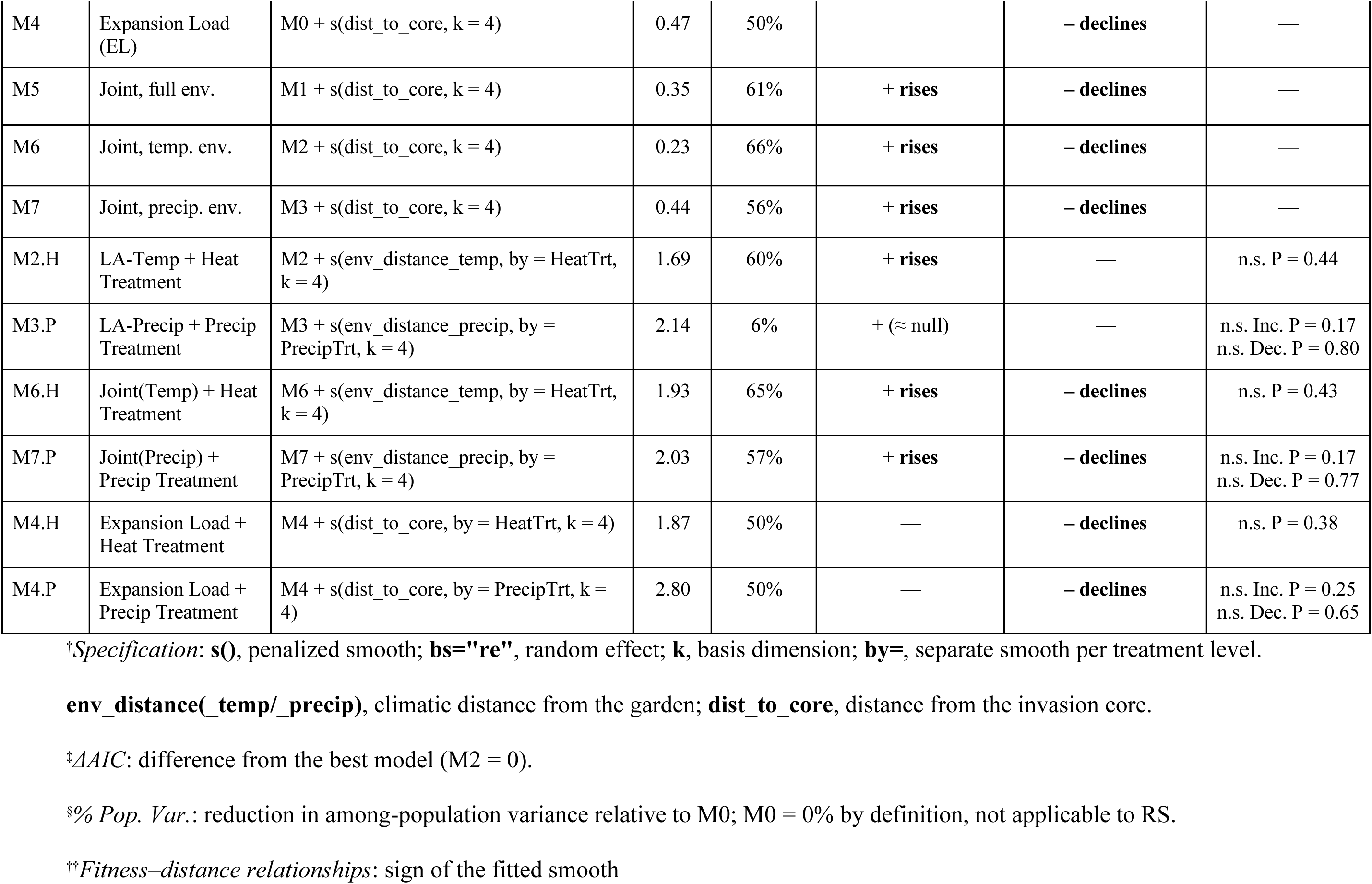
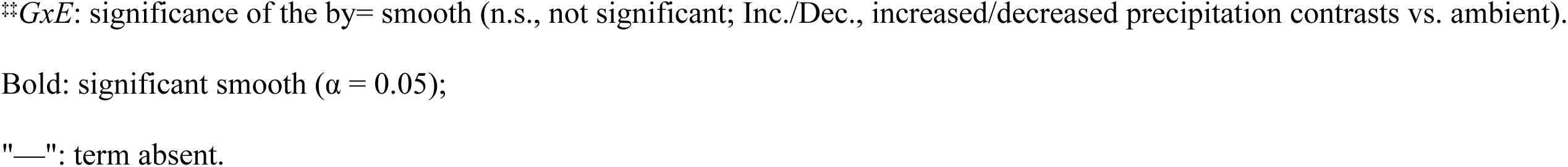
Fourteen candidate generalized additive mixed models (GAMs) testing for local adaptation and/or expansion load. Models tested whether among-population variation in composite fitness reflects local adaptation (fitness declining with source-to-garden environmental distance) or expansion load (fitness declining with distance from the invasion core). All models share the random-effects scaffold (RS); M0 adds the parametric climate treatments and is the reference model; M1–M7 add distance smooths; .H and .P suffixes add a treatment-specific smooth testing G × E.

| Model Name | Model Description (Hypothesis Tested) | Specification<sup>†</sup> | $\Delta AIC^{\ddagger}$ | % Pop. Var.<sup>§</sup> | Fitness - Env. Dist. relationship (Local Adaptation)<sup>††</sup> | Fitness - Dist. to Core relationship (Expansion Load)<sup>††</sup> | GxE relationship<sup>‡‡</sup> |
| --- | --- | --- | --- | --- | --- | --- | --- |
| RS | Random Effects Scaffold | $s(\text{Pop}, \text{bs}=\text{"re"}) + s(\text{Pop}, \text{HeatTrt}, \text{bs}=\text{"re"}) + s(\text{Pop}, \text{PrecipTrt}, \text{bs}=\text{"re"}) + s(\text{Block}, \text{bs}=\text{"re"})$ | — | — | — | — | — |
| M0 | Parametric Treatments | HeatTrt + PrecipTrt + HeatTrt:PrecipTrt + RS | 0.53 | 0% | — |  | — |
| M1 | Local Adaptation (LA) - full | M0 + $s(\text{env\_distance}, k = 4)$ | 0.78 | 56% | **+ rises** | | — |
| M2 | Local Adaptation - Temperature | M0 + $s(\text{env\_distance\_temp}, k = 4)$ | 0 | 60% | **+ rises** | | — |
| M3 | Local Adaptation - Precipitation | M0 + $s(\text{env\_distance\_precip}, k = 4)$ | 0.54 | 6% | **+ (≈ null)** | | — |
| M4 | Expansion Load (EL) | $M0 + s(\text{dist\_to\_core}, k = 4)$ | 0.47 | 50% | | – declines | — |
| M5 | Joint, full env. | $M1 + s(\text{dist\_to\_core}, k = 4)$ | 0.35 | 61% | + rises | – declines | — |
| M6 | Joint, temp. env. | $M2 + s(\text{dist\_to\_core}, k = 4)$ | 0.23 | 66% | + rises | – declines | — |
| M7 | Joint, precip. env. | $M3 + s(\text{dist\_to\_core}, k = 4)$ | 0.44 | 56% | + rises | – declines | — |
| M2.H | LA-Temp + Heat Treatment | $M2 + s(\text{env\_distance\_temp}, \text{by} = \text{HeatTrt}, k = 4)$ | 1.69 | 60% | + rises | — | n.s. P = 0.44 |
| M3.P | LA-Precip + Precip Treatment | $M3 + s(\text{env\_distance\_precip}, \text{by} = \text{PrecipTrt}, k = 4)$ | 2.14 | 6% | + (≈ null) | — | n.s. Inc. P = 0.17<br>n.s. Dec. P = 0.80 |
| M6.H | Joint(Temp) + Heat Treatment | $M6 + s(\text{env\_distance\_temp}, \text{by} = \text{HeatTrt}, k = 4)$ | 1.93 | 65% | + rises | – declines | n.s. P = 0.43 |
| M7.P | Joint(Precip) + Precip Treatment | $M7 + s(\text{env\_distance\_precip}, \text{by} = \text{PrecipTrt}, k = 4)$ | 2.03 | 57% | + rises | – declines | n.s. Inc. P = 0.17<br>n.s. Dec. P = 0.77 |
| M4.H | Expansion Load + Heat Treatment | $M4 + s(\text{dist\_to\_core}, \text{by} = \text{HeatTrt}, k = 4)$ | 1.87 | 50% | — | – declines | n.s. P = 0.38 |
| M4.P | Expansion Load + Precip Treatment | $M4 + s(\text{dist\_to\_core}, \text{by} = \text{PrecipTrt}, k = 4)$ | 2.80 | 50% | — | – declines | n.s. Inc. P = 0.25<br>n.s. Dec. P = 0.65 |

†*Specification: s()*, penalized smooth; **bs**="re", random effect; **k**, basis dimension; **by**=, separate smooth per treatment level.
**env\_distance(\_temp/\_precip)**, climatic distance from the garden; **dist\_to\_core**, distance from the invasion core.
‡*AAIC*: difference from the best model ( $M2 = 0$ ).
§% *Pop. Var.*: reduction in among-population variance relative to $M0$ ; $M0 = 0\%$ by definition, not applicable to RS.
††*Fitness–distance relationships*: sign of the fitted smooth
$\#GxE$ : significance of the by= smooth (n.s., not significant; Inc./Dec., increased/decreased precipitation contrasts vs. ambient).
Bold: significant smooth ( $\alpha = 0.05$ );
"—": term absent.

#### Population variation in fitness, main treatment effects, and G × E interactions

To test for fitness differences among population and climate treatments, we assessed our base model, M0, which contained only the shared random variables and the fixed linear treatment terms (Table 1). Because population was included as a random effect, we tested for variation in fitness among populations using a variance-component test of the GAM smooth population term, s(Pop, bs=“re”). We tested for the main effects of climate manipulations and their interaction using parametric F-tests. Similar to population, we tested for population-specific treatment responses (G × E) via variance-component tests of the GAM smooth population terms, and s(Pop, PrecipTrt, bs=“re”). For population-level analyses and figures, we extracted population-level estimated marginal means (EMM) of fitness from M0 (See Supplementary Material for details)

### Local adaptation versus expansion load

We compared seven additional candidate models against M0 to ask whether population-level variation in fitness was structured by environmental distance (consistent with local adaptation), geographic distance from the invasion core (consistent with expansion load), or both. Each candidate model added one or more penalized smooth terms to M0 (Table 1). We ranked candidate models by AIC and treated ΔAIC > 0.5 as substantial support for alternative models (Burnham & Anderson, 2002).

### Mechanistic factors contributing to expansion load

We evaluated the extent to which population attributes co-varied across the expansion gradient using Pearson correlations: distance from the invasion core, source-population size, Fixation Index (***F***), and experimental population fitness (M0 EMMs). We expected serial bottlenecks (expansion-front genetic drift) to yield a negative relationship between population size and distance from the invasion core and a positive correlation between ***F*** and distance from the invasion core. Both relationships were assessed using data from the statewide survey (N = 173 populations) and the subset that were source populations in the experiment (N = 13). In the statewide survey, fixation indices for seven individuals were considered outliers based on the IQR method and excluded from analyses.

### Fitness variation across populations in response to climate manipulation

To test our hypotheses concerning G x E, we fit six additional candidate models with distance smooths stratified by a matched treatment effect (Table 1). First, we tested for local adaptation to the manipulated climatic axes, where we expected the fitness-environmental distance relationship to vary by treatment if populations had locally adapted along that particular environmental axis (e.g., a significant s(env_distance_temp, by = HeatTrt); Table 1: M2.H, M3.P, M6.H, M7.P). Second, we tested for a signature of stress-amplified expansion load, where we expected edge populations would have greater reductions in fitness under stress resulting in greater negative slopes between fitness and distance from the core in heat or precipitation manipulations. We ranked these models using AIC along with the eight base models to determine which best fit our data. For each of the 14 models, we also report the proportion of among-population variance explained by the distance predictors, calculated as the proportional reduction in the population random-intercept variance relative to the null model M0 (100 x [1 - σ^2^_pop(model)_ / σ^2^_pop(M0)_]), together with the direction (sign) of each fitted distance smooth.

## Results

### Genetically based fitness differences across the expanding range

Source populations differed strongly in mean fitness (s(Pop): edf = 14.0, F = 29.8, P < 0.001; Fig. 2A & Fig S6), with approximately 2.7× greater fitness for the highest (COOK003) than lowest performing population (BIG002) (range: 1.9 ± 0.18 to 5.1 ± 0.27; mean ± SE). Block also contributed appreciable variance (s(block): edf = 6.4, F = 1.5, P = 0.002). The base model (M0) including the random effect of population explained 43% of the variation in fitness. Fixed treatment effects in M0 accounted for only 2.8% of the variation in fitness.

**Figure 2.**
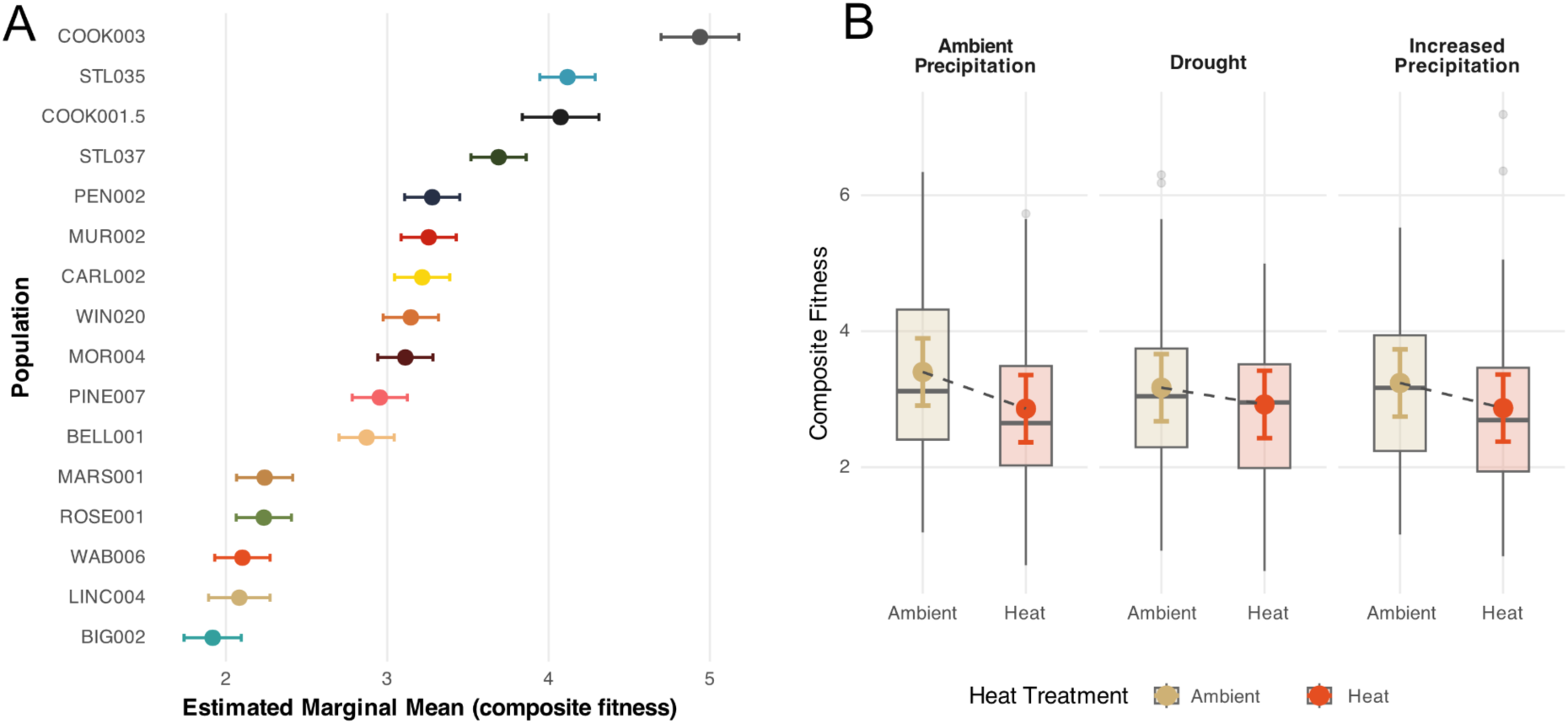
A) Marginal mean population fitness of the 16 experimental source populations from GAM model M0. Composite fitness estimates (± SE) are colored by population and ordered by value. B) Fixed experimental treatment effects on composite fitness from GAM model M0. Composite fitness estimates are grouped by the precipitation treatment x heat treatment interaction: (left) Ambient precipitation, (center) 30% decreased precipitation (drought) using event-based rain-out shelters, (right) 30% precipitation additions. Within each precipitation treatment group, box plots and estimated mean values (± SE) for ambient temperatures are shown in muted gold and heat treatments are shown in red. **Alt. Text**: Graphs showing population mean fitness with subpanels labeled A and B showing mean fitness for each population in the experiment and mean fitness for the climate manipulations.

### Warming, but not altered precipitation, reduced fitness

Elevated temperature reduced fitness by 11.3% (F₁ = 12.1, *P* < 0.001; ambient: 3.3 ± 0.08, elevated: 2.9 ± 0.08; Fig. 2B). However, we found no effect of precipitation (F₂ = 0.2, *P* = 0.84) or a temperature × precipitation interaction (F₂ = 1.0, *P* = 0.39). We also did not find evidence for G × E interactions for temperature (s(Pop, temperature): F = 0.5, *P* = 0.17) or precipitation (F ≈ 0, *P* = 0.58). Therefore, the warming-induced fitness decline was largely shared across source populations.

### Mean fitness declined from the invasion core to expanding range margins

The four best-supported models were within 0.5 ΔAIC of each other (M2 temperature-distance only; M6 joint temperature-distance + core-distance; M5 joint full-environment-distance + core-distance; M7 joint precipitation-distance + core-distance; Table 1; Fig. S7), and a core-distance-only model was within 0.6 ΔAIC (M4; Table 1; Fig. 3B). Within-model smooth tests identified temperature distance and distance to core as the strongest individual predictors of among-population fitness variation; precipitation distance was never a significant predictor (Figure 3; Table S4).

**Figure 3.**
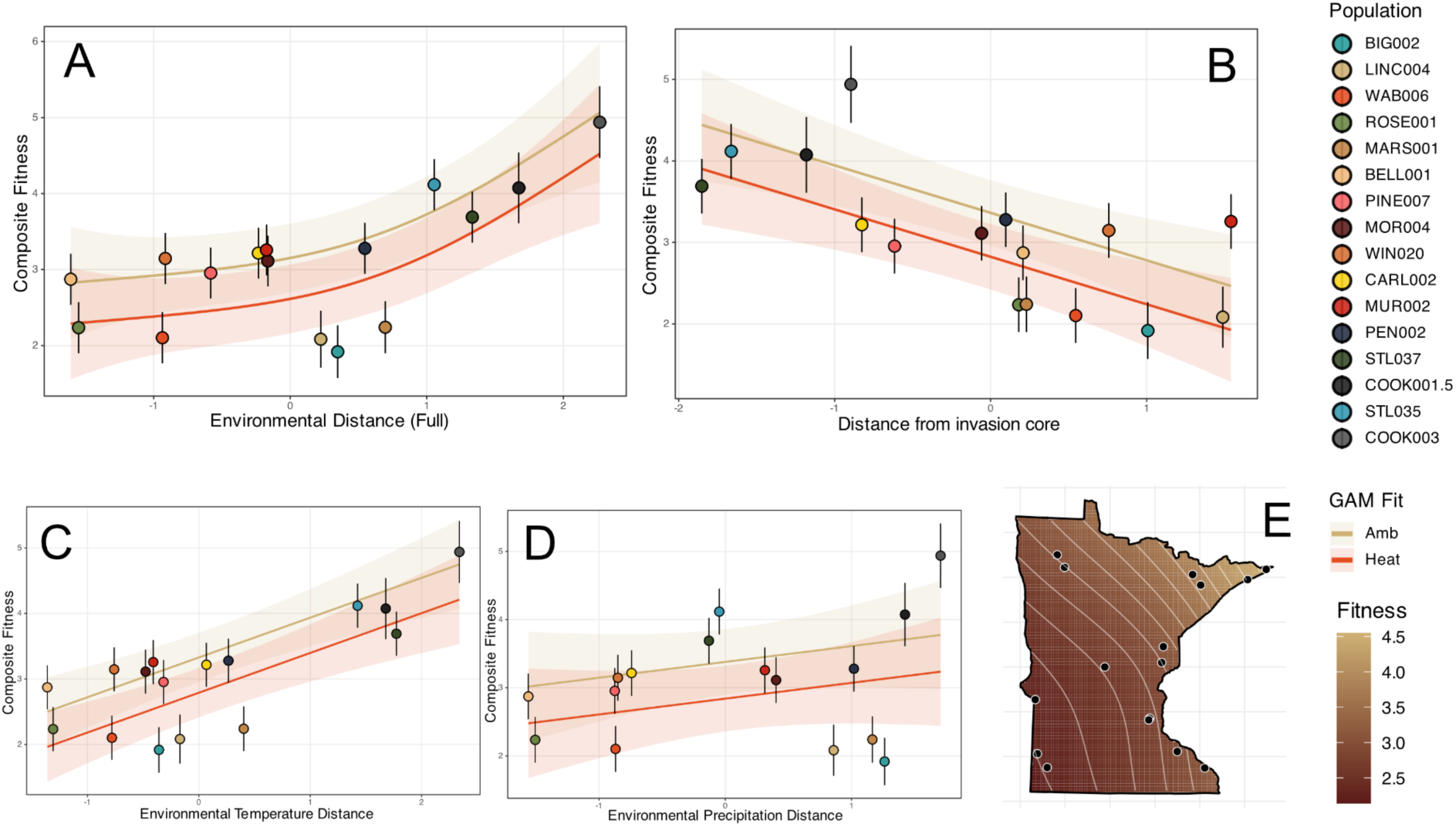
Correlation from single distance predictor models between population mean composite fitness and A) Full environmental distance (M1: local adaptation test) B) distance from the invasion core (M4: expansion load test), C) Environmental temperature distance (M2: thermal local adaptation test), and D) Environmental precipitation distance (M3: precipitation local adaptation test). Lines with confidence intervals represent GAM fits from the specified model grouped by heated (red) or ambient (muted gold) heat treatments. Estimated population mean fitness (± SE) plotted and colored by population. E) Spatial interpolation of mean population fitness across Minnesota fit using a two-dimensional smooth GAM with a thin-plate regression spline on UTM-projected source coordinates: gam(emm ∼ s(Easting, Northing, k = 4)). **Alt. Text**: Graphs and map in panels labeled A-E showing how fitness is related to different measures of environmental distance or distance from the center of the invasion core.

Temperature distance and distance to core were confounded in our dataset (raw data: r = −0.70; M6; joint model worst-case concurvity = 0.77). In our single-predictor models, M2 (temperature distance; Fig 3C) and M4 (distance to core; Fig. 3B), explained similar proportions of among-population variance (60% v. 50%) (Table 1). However, only our expansion load hypothesis was supported based on predictions (Table 1). Fitness declined from the northeastern core toward both the expanding southern and western range margins (Fig. 3E). Conversely, the fitness-temperature distance relationship was always positive (Table 1), suggesting local maladaptation rather than local adaptation (Fig. 3C).

### Mechanistic signals of expansion load

We predicted that increased expansion load would result in edge populations that are smaller and have greater homozygosity (higher ***F***) than core populations. Across the statewide survey, population size declined (r = −0.50, *P* < 0.001; Fig. 4B) and fixation index (***F***) increased (r = 0.26, *P* < 0.001; Fig. 4A) with distance from the invasion core; population size and ***F*** were uncorrelated (r = −0.01, *P* = 0.87) (Table S5). Similarly, for experimental populations, population size declined (r = −0.73, *P* = 0.005; Fig. 4D) and ***F*** increased with distance from the core (r = 0.81, *P* = 0.001; Fig. 4C). The exclusion of outliers did not impact results (See Fig. S8 & Table S6)

**Figure 4.**
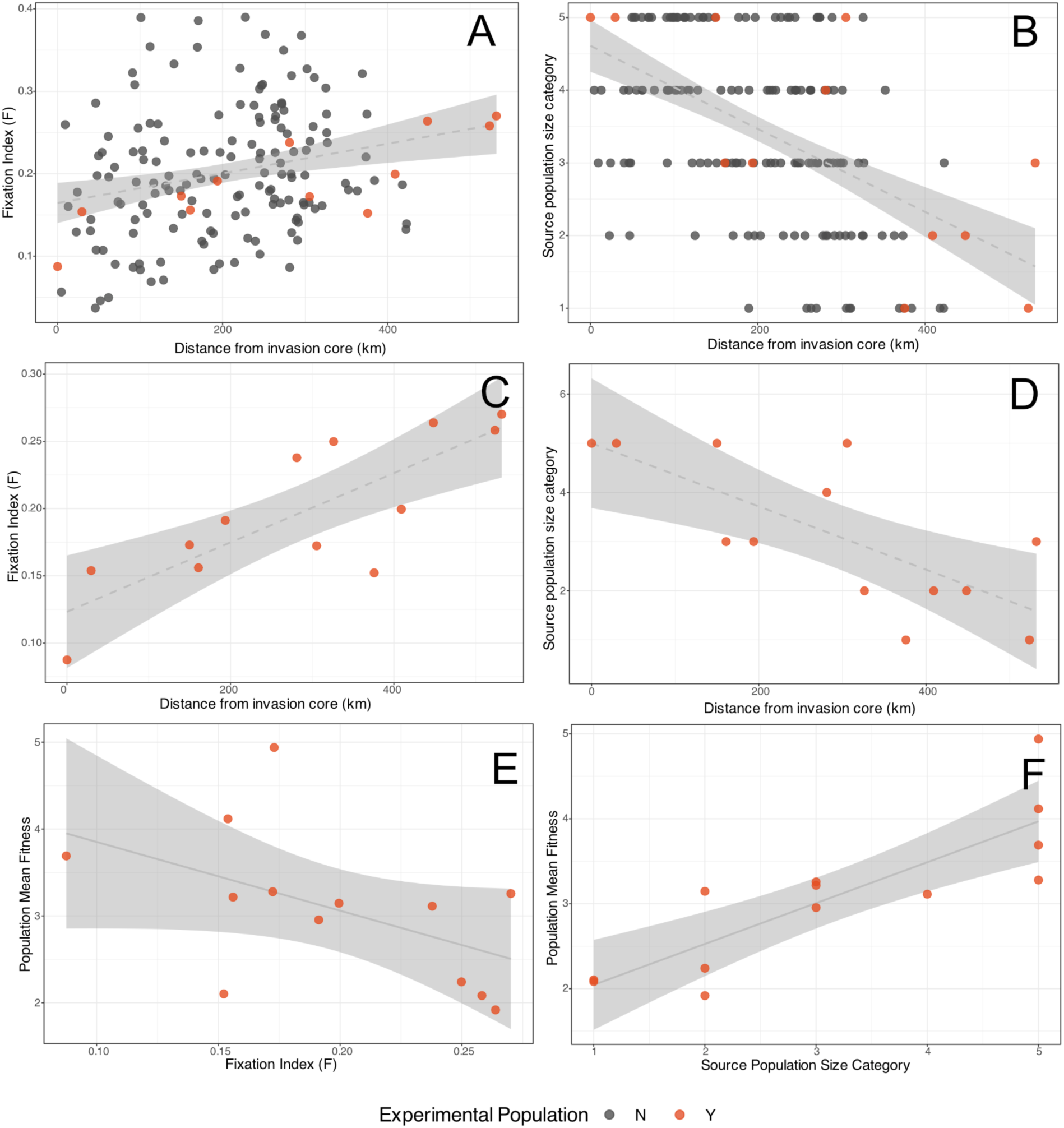
Potential mechanisms underlying expansion load. (A-D) Common tansy population relationships between distance from the invasion core and population Fixation Index (***F***) for A) all populations and C) only experimental populations OR Estimated Source Population Size for B) all populations and D) only experimental populations. (E-F) Relationship between estimated mean fitness of experimental populations and E) Fixation Index (***F***) and F) source population size. Experimental populations shown with orange points; all other surveyed populations shown in gray. **Alt Text**: Graphs of data on potential mechanisms of expansion load with subpanels labeled A-F. Panels show how either fixation index or population size was related to distance from the invasion core or mean population fitness.

Population-level fitness (M0 EMMs) was strongly predicted by source population size (r = 0.84, *P* < 0.001; Fig. 4F). The correlation between mean population fitness and ***F*** was negative but not quite significant (r = −0.51, *P* = 0.08; Fig. 4E).

### No evidence for G × E along temperature, precipitation, or invasion gradients

To further investigate the processes driving population differentiation, we tested two specific G × E predictions. First, we tested for a *population source climate* × *treatment* interaction indicative of local adaptation, expecting that a population’s fitness response to climate manipulation would change the fitness-environmental distance relationship and depend predictably on its climate of origin. Second, we tested for stress-amplified expansion load, predicting that the fitness ∼ distance-to-core slope would steepen under environmental stress. None of the G × E tests for local adaptation were significant, suggesting that climate of origin did not influence population responses to climate manipulations (Table 1). Similarly, there was no indication of stress-amplified load. Populations sourced near the expanding range margins did not respond differently to manipulations (Table 1). Additionally, G × E terms did not improve model performance as all models were ranked lower than simpler models that excluded these terms in the AIC table (ΔAIC ≥ 1.7; Table 1).

## Discussion

Disentangling selective and stochastic forces shaping fitness during range expansion clarifies their influence on range limits (Holtz et al., 2025; Miller et al., 2020; Polechová & Barton, 2015) and helps predict population responses to environmental change (Abeli et al., 2014; Bridle & Hoffmann, 2022; Gilbert et al., 2018; Sexton et al., 2009). We conducted a manipulative field experiment and asked whether fitness across the invaded range is better explained by local adaptation or expansion load. We show that populations have diverged strongly in fitness over the ca. 150 year period of range expansion. Population mean fitness declined from the long-established northeastern core toward the southern and western range margins. We found no signal of local adaptation along environmental gradients; rather, higher fitness was realized in populations sourced from more dissimilar environments. Instead, the geographic fitness decline tracked independent demographic (declining population size) and genetic (excess homozygosity) signatures of serial founding events. Overall, our results suggest that expansion load is an important factor influencing fitness at the range margins of this young invasion.

Mean fitness (composite of vegetative and reproductive biomass) varied roughly 2.7-fold and reproductive biomass varied roughly 12-fold across source populations. Although these differences are pronounced, it should be noted that common tansy is a perennial and populations may differ in their allocation to growth vs. reproduction and individual lifespan. However, our experiment tracked fitness over two seasons and included marginal populations that spanned different environmental gradients (i.e. western and southern) with no evidence of local adaptation from any fitness metric.

Two non-exclusive hypotheses predict the rapid divergence in fitness across the invasion chronosequence: local adaptation along climatic gradients and expansion load at range margins. While fitness variation was related to temperature distance, this result reflected local maladaptation rather than adaptation (Kawecki & Ebert, 2004). While maladaptation could arise due to mismatch between historical and current climate (e.g., Anderson & Wadgymar, 2019; Gorton et al., 2022), the best-performing populations in our study originated from cooler conditions - despite warmer-than-average experimental years. Distance to core, by contrast, has a clear mechanistic underpinning – serial founding events and the genetic load it generates. As we discuss below, evidence for expansion load was independently corroborated by demographic and genomic datasets. Because the common garden sits near the southern expanding margin rather than in the northeastern core, temperature distance and distance-to-core were negatively correlated, which allowed us to distinguish between local adaptation and expansion load hypotheses.

If genetic load drives the geographic pattern of fitness decline, the serial founder events that produced it should leave demographic and genomic footprints. Indeed, we found that marginal populations had smaller sizes and higher homozygosity genome-wide. Population size fell with distance from the core and was the strongest correlate of population fitness. Homozygosity rose toward the edge (Fig 4A & 4C) and had a negative but non-significant relationship to fitness (r = −0.51, P = 0.08). While these demographic and genomic signatures implicate genetic drift as an important evolutionary process at the expansion front, it is difficult to know the exact causal relationships between population size and fitness (small population size leads to genetic load and reduced fitness versus the reverse). In future work, more genomic analyses could illuminate demographic history of the expansion in more detail and also assess molecular evidence of deleterious load (González-Martínez et al., 2017; Rougemont et al., 2023; Willi et al., 2018). Crossing experiments within and between core and marginal populations could be used to precisely quantify the load incurred by deleterious recessive mutations (sensu Perrier et al., 2020, 2022) and reveal the extent to which gene flow could relieve expansion load and rescue fitness at and beyond the expanding range edge.

Expansion load is well grounded in theory and documented at native range edges (Koski et al., 2019; Peischl et al., 2015; Perrier et al., 2020). However, it is rarely invoked for contemporary invasions, where the focus has been adaptation and the purging of genetic load (Colautti & Lau, 2015; Estoup et al., 2016; van Kleunen et al., 2018). Our results suggest that rapid fitness divergence was driven by stochastic processes that reduce fitness, not by adaptation to the new environments it crossed. Whether load generally dominates active expansions is an open question and one that reaches beyond non-native invasions. In cases where native species exhibit range shifts due to climate change, do their advancing edges carry expansion load that current range-shift models overlook? The answer will depend on historical population size, the supply and recessivity of deleterious mutations, the severity of founder events, and how quickly selection purges load behind the front (Gilbert et al., 2018; Peischl et al., 2015).

## Data and Code Availability Statement

All data and code is archived at the Data Repository of the University of Minnesota (DRUM) (doi: xxxx).

## Author Contributions

RBR and DAM planned and conducted the experiment. RBR and JWB conducted statistical analysis. All authors contributed to the drafting and editing of the manuscript.

## Acknowledgements

We thank R. Venette and A. Morey for thoughtful comments on the results of our preliminary analyses. We thank the Minnesota Department of Agriculture and the Minnesota Department of Natural Resources, in particular T. Cortilet and L. Van Riper, for help with permitting and identification of populations. We thank A. Scobbie, J. Leonard and the UMN MAES St. Paul Campus Research Facilities for assistance with setting up our experiment. We thank T. Lake, D. Schoenecker, B. Greene, I. Olsen for field assistance and members of the Moeller Lab for help during harvest.

## References

Abeli, T., Gentili, R., Mondoni, A., Orsenigo, S., & Rossi, G. (2014). Effects of marginality on plant population performance. Journal of Biogeography, 41(2), 239–249.

Anderson, J. T., & Wadgymar, S. M. (2019). Climate change disrupts local adaptation and favours upslope migration. Ecology Letters, 23(1), 181–192.

Angert, A. L., Bontrager, M. G., & Ågren, J. (2020). What Do We Really Know About Adaptation at Range Edges? Annual Review of Ecology, Evolution, and Systematics, 51(1), 341–361.

Baker, H. G., & Stebbins, G. L. (eds). (1965). The genetics of colonizing species. Academic Press, New York.

Becker, R. A., Wilks, A. R., Brownrigg, R., Minka, T. P., & Deckmyn, A. (2025). maps: Draw Geographical Maps. https://CRAN.R-project.org/package=maps

Bosshard, L., Dupanloup, I., Tenaillon, O., Bruggmann, R., Ackermann, M., Peischl, S., & Excoffier, L. (2017). Accumulation of deleterious mutations during bacterial range expansions. Genetics, 207(2), 669–684.

Bridle, J., & Hoffmann, A. (2022). Understanding the biology of species’ ranges: when and how does evolution change the rules of ecological engagement? Philosophical Transactions of the Royal Society of London. Series B, Biological Sciences, 377(1848), 20210027.

Briscoe Runquist, R., & Moeller, D. A. (2024). Isolation by environment and its consequences for range shifts with global change: Landscape genomics of the invasive plant common tansy. Molecular Ecology, 33(16), e17462.

Brun, P., Zimmermann, N. E., Hari, C., Pellissier, L., & Karger, D. N. (9/2025). CHELSA-bioclim [Dataset]. In Chelsa-BIOCLIM+ A novel set of global climate-related predictors at kilometre-resolution. 10.16904/envidat.332

Burnham, K. P., & Anderson, D. R. (2002). Model selection and multimodel inference: A practical information-theoretic approach (K. P. Burnham & D. R. Anderson, Eds.; 2nd ed.) [PDF]. Springer.

Chen, L., & Ford, T. W. (2023). Future changes in the transitions of monthly-to-seasonal precipitation extremes over the Midwest in Coupled Model Intercomparison Project Phase 6 models. International Journal of Climatology: A Journal of the Royal Meteorological Society, 43(1), 255–274.

Cheptou, P. O., Berger, A., Blanchard, A., Collin, C., & Escarre, J. (2000). The effect of drought stress on inbreeding depression in four populations of the Mediterranean outcrossing plant Crepis sancta (Asteraceae). Heredity, 85 Pt 3(3), 294–302.

Clements, D. R., & Jones, V. L. (2021). Rapid Evolution of Invasive Weeds Under Climate Change: Present Evidence and Future Research Needs. Frontiers in Agronomy, 3. 10.3389/fagro.2021.664034

Colautti, R. I., & Lau, J. A. (2015). Contemporary evolution during invasion: evidence for differentiation, natural selection, and local adaptation. Molecular Ecology, 24(9), 1999–2017.

Colautti, R. I., Maron, J. L., & Barrett, S. C. H. (2009). Common garden comparisons of native and introduced plant populations: latitudinal clines can obscure evolutionary inferences. Evolutionary Applications, 2(2), 187–199.

Dlugosch, K. M., & Parker, I. M. (2008). Founding events in species invasions: genetic variation, adaptive evolution, and the role of multiple introductions. Molecular Ecology, 17(1), 431–449.

Estoup, A., Ravigné, V., Hufbauer, R., Vitalis, R., Gautier, M., & Facon, B. (2016). Is there a genetic paradox of biological invasion? Annual Review of Ecology, Evolution, and Systematics, 47(1), 51–72.

Excoffier, L., Foll, M., & Petit, R. J. (2009). Genetic consequences of range expansions. Annual Review of Ecology, Evolution, and Systematics, 40(1), 481–501.

Excoffier, L., & Ray, N. (2008). Surfing during population expansions promotes genetic revolutions and structuration. Trends in Ecology & Evolution, 23(7), 347–351.

Ford, T. W., Chen, L., & Schoof, J. T. (2021). Variability and transitions in precipitation extremes in the Midwest United States. Journal of Hydrometeorology, 22(3), 533–545.

Fox, C. W., & Reed, D. H. (2011). Inbreeding depression increases with environmental stress: an experimental study and meta-analysis: Inbreeding load increases with stress. Evolution; International Journal of Organic Evolution, 65(1), 246–258.

Frank, D., & Klotz, S. (1988). Biologisch-ökologische Daten zur Flora der DDR. 2nd ed. Martin-Luther University Halle-Wittenberg, Halle (Saale) DE

Gilbert, K. J., Peischl, S., & Excoffier, L. (2018). Mutation load dynamics during environmentally-driven range shifts. PLoS Genetics, 14(9), e1007450.

Gilbert, K. J., Sharp, N. P., Angert, A. L., Conte, G. L., Draghi, J. A., Guillaume, F., Hargreaves, A. L., Matthey-Doret, R., & Whitlock, M. C. (2017). Local adaptation interacts with expansion load during range expansion: Maladaptation reduces expansion load. The American Naturalist, 189(4), 368–380.

González-Martínez, S. C., Ridout, K., & Pannell, J. R. (2017). Range expansion compromises adaptive evolution in an outcrossing plant. Current Biology, 27(16), 2544–2551.e4.

Gorton, A. J., Benning, J. W., Tiffin, P., & Moeller, D. A. (2022). The spatial scale of adaptation in a native annual plant and its implications for responses to climate change. Evolution; International Journal of Organic Evolution, 76(12), 2916–2929.

Henn, B. M., Botigué, L. R., Peischl, S., Dupanloup, I., Lipatov, M., Maples, B. K., Martin, A. R., Musharoff, S., Cann, H., Snyder, M. P., Excoffier, L., Kidd, J. M., & Bustamante, C. D. (2016). Distance from sub-Saharan Africa predicts mutational load in diverse human genomes. Proceedings of the National Academy of Sciences of the United States of America, 113(4), E440–9.

Hewitt, G. (2000). The genetic legacy of the Quaternary ice ages. Nature, 405(6789), 907–913.

Hijmans, R. J. (2026). geosphere: Spherical Trigonometry. https://github.com/rspatial/geosphere

Hodgins, K. A., Battlay, P., & Bock, D. G. (2025). The genomic secrets of invasive plants. The New Phytologist, 245(5): 1846–1863.

Holt, R. D. (2003). On the evolutionary ecology of species’ ranges. Evolutionary Ecology Research, 5(2), 159–178.

Holtz, S., Hudson, A., Tittes, S., & Weiss-Lehman, C. (2025). Genetic consequences of gene surfing and their relationship to fitness. Evolution; International Journal of Organic Evolution, 79(10), 2208–2218.

Jacobs, J. (2008). Ecology and Management of Common Tansy (Tanacetum vulgare L.). USDA Natural Resources Conservation Service.

Kawecki, T. J., & Ebert, D. (2004). Conceptual issues in local adaptation. Ecology Letters, 7(12), 1225–1241.

Kirkpatrick, M., & Barton, N. H. (1997). Evolution of a species’ range. The American Naturalist, 150(1), 1–23.

Klopfstein, S., Currat, M., & Excoffier, L. (2006). The Fate of Mutations Surfing on the Wave of a Range Expansion. Molecular Biology and Evolution, 23(3), 482–490.

Koski, M. H., Layman, N. C., Prior, C. J., Busch, J. W., & Galloway, L. F. (2019). Selfing ability and drift load evolve with range expansion. Evolution Letters, 3(5), 500–512.

Kunkel, K.E, Frankson, R., Runkle, J., Champion, S. M., Stevens, L. E., Easterling, D. R., Stewart, B. C., McCarrick, A., & Lemery (Eds.), C. (2022). State Climate Summaries for the United States 2022. NOAA Technical Report NESDIS 150. https://statesummaries.ncics.org/chapter/mn

Lake, T. A., Briscoe Runquist, R. D., & Moeller, D. A. (2020). Predicting range expansion of invasive species: Pitfalls and best practices for obtaining biologically realistic projections. Diversity and Distributions, 26(12), 1767–1779.

LeCain, R., & Sheley, R. (2014). Common Tansy: A self-learning resource from MSU extension (No. MT199911AG). Montana State University. https://www.montana.edu/extension/invasiveplants/documents/mt_noxious_weeds/common_tansy.pdf

Liess, S., Twine, T. E., Snyder, P. K., Hutchison, W. D., Konar-Steenberg, G., Keeler, B. L., & Brauman, K. A. (2022). High-Resolution Climate Projections Over Minnesota for the 21st Century. Earth and Space, 9(3), e2021EA001893.

Lokki, J., Sorsa, M., Forsén, K., & Schantz, M. V. (1973). Genetics of monoterpenes in Chrysanthemum vulgare: I. Genetic control and inheritance of some of the most common chemotypes. Hereditas, 74(2), 225–232.

Mack, R. N. (2003). Plant Naturalizations and Invasions in the Eastern United States: 1634-1860. Annals of the Missouri Botanical Garden, 90(1), 77–90.

Miller, T. E. X., Angert, A. L., Brown, C. D., Lee-Yaw, J. A., Lewis, M., Lutscher, F., Marculis, N. G., Melbourne, B. A., Shaw, A. K., Szűcs, M., Tabares, O., Usui, T., Weiss-Lehman, C., & Williams, J. L. (2020). Eco-evolutionary dynamics of range expansion. Ecology, 101(10), e03139.

Minnesota Department of Natural Resources. (2021-2022). U of M St. Paul Campus Climate Observatory. https://www.dnr.state.mn.us/climate/climate_monitor/climate_observatory.html

Mitich, L. W. (1992). Tansy. Weed Technology: A Journal of the Weed Science Society of America, 6(1), 242–244.

Oduor, A. M. O., Leimu, R., & van Kleunen, M. (2016). Invasive plant species are locally adapted just as frequently and at least as strongly as native plant species. The Journal of Ecology, 104(4), 957–968.

Peischl, S., Dupanloup, I., Foucal, A., Jomphe, M., Bruat, V., Grenier, J.-C., Gouy, A., Gilbert, K. J., Gbeha, E., Bosshard, L., Hip-Ki, E., Agbessi, M., Hodgkinson, A., Vézina, H., Awadalla, P., & Excoffier, L. (2018). Relaxed selection during a recent human expansion. Genetics, 208(2), 763–777.

Peischl, S., Dupanloup, I., Kirkpatrick, M., & Excoffier, L. (2013). On the accumulation of deleterious mutations during range expansions. Molecular Ecology, 22(24), 5972–5982.

Peischl, S., & Excoffier, L. (2015). Expansion load: recessive mutations and the role of standing genetic variation. Molecular Ecology, 24(9), 2084–2094.

Peischl, S., & Excoffier, L. (2016). Expansion load: Recessive mutations and the role of standing genetic variation. In Invasion Genetics (pp. 218–231). John Wiley & Sons, Ltd.

Peischl, S., Kirkpatrick, M., & Excoffier, L. (2015). Expansion load and the evolutionary dynamics of a species range. The American Naturalist, 185(4), E81–93.

Perrier, A., Sánchez-Castro, D., & Willi, Y. (2020). Expressed mutational load increases toward the edge of a species’ geographic range. Evolution; International Journal of Organic Evolution, 74(8), 1711–1723.

Perrier, A., Sánchez-Castro, D., & Willi, Y. (2022). Environment dependence of the expression of mutational load and species’ range limits. Journal of Evolutionary Biology, 35(5), 731–741.

Peschel, A. R., Flint, S. A., May, G., & Shaw, R. G. (2025). Genetic divergence in population mean fitness is weakly associated with environmental and geographic distance in four prairie perennial forbs. Evolution Letters, 9(5), 522–532.

Polechová, J. (2018). Is the sky the limit? On the expansion threshold of a species’ range. PLoS Biology, 16(6), e2005372.

Polechová, J., & Barton, N. H. (2015). Limits to adaptation along environmental gradients. Proceedings of the National Academy of Sciences of the United States of America, 112(20), 6401–6406.

Prach, K., & Wade, P. M. (1992). Population characteristics of expansive perennial herbs. Preslia. https://agris.fao.org/agris-search/search.do?recordID=CS9201864

Purcell, S. (2019). PLINK v. 1.9. http://pngu.mgh.harvard.edu/purcell/plink/

Purcell, S., Neale, B., Todd-Brown, K., Thomas, L., Ferreira, M. A. R., Bender, D., Maller, J., Sklar, P., de Bakker, P. I. W., Daly, M. J., & Sham, P. C. (2007). PLINK: a tool set for whole-genome association and population-based linkage analyses. American Journal of Human Genetics, 81(3), 559–575.

Roberts, T. (1878). *Tanacetum vulgare* (University of Minnesota). Bell Atlas Detailed Collection Record Information. https://bellatlas.umn.edu/collections/individual/index.php?occid=158375

Rougemont, Q., Leroy, T., Rondeau, E. B., Koop, B., & Bernatchez, L. (2023). Allele surfing causes maladaptation in a Pacific salmon of conservation concern. PLoS Genetics, 19(9), e1010918.

Schrieber, K., & Lachmuth, S. (2017). The Genetic Paradox of Invasions revisited: the potential role of inbreeding × environment interactions in invasion success: The Genetic Paradox revisited. Biological Reviews of the Cambridge Philosophical Society, 92(2), 939–952.

Sexton, J. P., McIntyre, P. J., Angert, A. L., & Rice, K. J. (2009). Evolution and Ecology of Species Range Limits. Annual Review of Ecology, Evolution, and Systematics, 40, 415–436.

Uller, T., & Leimu, R. (2011). Founder events predict changes in genetic diversity during human-mediated range expansions. Global Change Biology, 17(11), 3478–3485.

UMN Climate Observatory. (2021-2022). U of M St. Paul Campus Climate Observatory Minnesota Department of Natural Resources. https://www.dnr.state.mn.us/climate/climate_monitor/climate_observatory.html

van Kleunen, M., Bossdorf, O., & Dawson, W. (2018). The Ecology and Evolution of Alien Plants. Annual Review of Ecology, Evolution, and Systematics, 49(1), 25–47.

Weiss-Lehman, C., Hufbauer, R. A., & Melbourne, B. A. (2017). Rapid trait evolution drives increased speed and variance in experimental range expansions. Nature Communications, 8, 14303.

White, D. J. (1997). Tanacetum vulgare L: weed potential, biology, response to herbivory, and prospects for classical biological control in Alberta. Ph. D. Dissertation Thesis, University of Alberta.

Wickham, H. (2016). ggplot2: Elegant Graphics for Data Analysis. Springer-Verlag New York. https://ggplot2.tidyverse.org

Wickham, H., Averick, M., Bryan, J., Chang, W., McGowan, L., François, R., Grolemund, G., Hayes, A., Henry, L., Hester, J., Kuhn, M., Pedersen, T., Miller, E., Bache, S., Müller, K., Ooms, J., Robinson, D., Seidel, D., Spinu, V., … Yutani, H. (2019). Welcome to the tidyverse. Journal of Open Source Software, 4(43), 1686.

Willi, Y., Fracassetti, M., Zoller, S., & Van Buskirk, J. (2018). Accumulation of Mutational Load at the Edges of a Species Range. Molecular Biology and Evolution, 35(4), 781–791.

Wolf, V. C., Gassmann, A., Clasen, B. M., Smith, A. G., & Müller, C. (2012). Genetic and chemical variation of Tanacetum vulgare in plants of native and invasive origin. Biological Control: Theory and Applications in Pest Management, 61(3), 240–245.

Wood, S. N. (2017). Generalized additive models: An introduction with R. Chapman and Hall/CRC.

Yang, J., Lee, S. H., Goddard, M. E., & Visscher, P. M. (2011). GCTA: a tool for genome-wide complex trait analysis. American Journal of Human Genetics, 88(1), 76–82.

