## Supplementary Material for "Expansion load reduces fitness at the range margin of an invasive plant"

**Supplementary Methods** **3**
Table S1. Minnesota Common Tansy Population Survey 3
Experimental Design Details 7
Figure S1. Experimental Plot Layout 7
Planting 8
Climate manipulations 8
Figure S2. OTC Temperature manipulations 9
Figure S3. Precipitation manipulations 10
Detailed fitness PCs 11
Figure S4. Composite fitness 11
Climate Variables 12
Table S2. Chelsea Climate Variables 12
Figure S5. Chelsa ClimateVariables 13
Table S3. Distance Correlations 14
Estimates of Population Mean Fitness 14
**Supplementary Results** **15**
Figure S6. Observed population mean fitness and fitness variation 15
Table S4. GAM effects table 16
Figure S7. Best joint GAM relationships 19
Table S5. Correlation among expansion load factors excluding outliers 19
Table S6. Correlations among expansion load factors including outliers 20
Figure S8. Components of expansion load with outliers 20

### Supplementary Methods

#### Table S1. Minnesota Common Tansy Population Survey

Population information of surveyed Minnesota common tansy populations. Population Sizes are based on field surveys done at time of tissue or seed collection. The numbers correspond to the following population census size estimate: 1 = 1-10 ind., 2 = 10-100 ind., 3 = 100-1000 ind., 4 = 1000-10000 ind., 5 = 10000+ ind. Bolded populations were those included in the common garden experiment. Italicized and starred (*) populations were considered F-statistic outliers using the IQR method and were excluded from analyses using F-statistics.

| Population | Latitude | Longitude | Pop. Size Cat. | F | Population | Latitude | Longitude | Pop. Size Cat. | F |
|---|---|---|---|---|---|---|---|---|---|
| **BELL001** | **44.988** | **-93.174** | **2** | | ITA013 | 47.226 | -93.497 | 5 | 0.1214 |
| **BIG002** | **45.345** | **-96.401** | **2** | **0.2309** | ITA014 | 47.088 | -93.197 | 5 | 0.1099 |
| **CARL002** | **46.459** | **-92.755** | **3** | **0.0641** | ITA015 | 47.204 | -93.763 | 4 | -0.0076 |
| **COOK001.5** | **47.774** | **-90.243** | **3** | | ITA017 | 47.293 | -93.703 | 3 | 0.1735 |
| **COOK003** | **47.965** | **-89.678** | **5** | **0.1092** | ITA018 | 47.089 | -93.196 | 5 | 0.0864 |
| **LINC004** | **44.262** | **-96.279** | **1** | **0.0921** | ITA019 | 47.396 | -93.086 | 5 | -0.0793 |
| **MOR004** | **46.042** | **-94.443** | **4** | **0.1163** | KOO003 | 48.646 | -94.166 | 3 | 0.0563 |
| **MUR002** | **43.992** | **-95.990** | **3** | **0.1186** | KOO004 | 48.629 | -93.863 | 3 | 0.2819 |
| **PEN002** | **48.020** | **-95.700** | **5** | **0.1780** | KOO005 | 48.519 | -93.619 | 3 | 0.1716 |
| **PINE007** | **46.140** | **-92.804** | **3** | **0.1087** | KOO006 | 48.280 | -93.737 | 3 | 0.2401 |
| **ROSE001** | **45.023** | **-93.146** | **2** | | KOO007 | 48.191 | -93.801 | 2 | 0.1810 |
| **STL035** | **47.902** | **-91.877** | **5** | **0.0776** | KOO008 | 47.964 | -94.145 | 4 | 0.0434 |
| **STL037** | **47.689** | **-91.640** | **5** | **-0.0454** | KOO009 | 47.856 | -94.345 | 2 | 0.1853 |

| | | | | | | | | | |
|---|---|---|---|---|---|---|---|---|---|
| **WAB006** | **44.352** | **-92.376** | **1** | **0.0205** | LAKE001 | 47.018 | -91.665 | 5 | -0.1751 |
| **WIN020** | **44.014** | **-91.616** | **2** | **0.0396** | LAKE004_24 | 47.272 | -91.328 | 5 | 0.0755 |
| **AIT008* | *46.987* | *-93.666* | *5* | *-0.2702* | LAKE004_25 | 47.414 | -91.238 | 4 | -0.0595 |
| **LAKE004_1* | *47.414* | *-91.238* | *4* | *0.5289* | LAKE006 | 47.730 | -91.645 | 4 | -0.0411 |
| **STL023* | *47.861* | *-92.718* | *4* | *0.5121* | LAKE008 | 47.473 | -91.645 | 3 | 0.0945 |
| AIT001 | 46.521 | -93.580 | 2 | 0.0115 | LAKE009 | 47.346 | -91.466 | 3 | 0.1064 |
| AIT002 | 46.986 | -93.714 | 3 | -0.0022 | LAKE010 | 47.325 | -91.331 | 5 | 0.0950 |
| AIT003 | 46.735 | -93.272 | 3 | 0.0446 | LAKE011 | 47.242 | -91.442 | 5 | -0.0182 |
| AIT005 | 46.926 | -93.294 | 3 | -0.0242 | LAKE012 | 47.207 | -91.570 | 5 | 0.0972 |
| AIT006 | 46.707 | -93.477 | 5 | -0.1498 | LAKE013 | 47.031 | -91.667 | 5 | 0.1295 |
| AIT007 | 46.826 | -93.612 | 5 | 0.1350 | LAKE014 | 47.258 | -91.656 | 3 | 0.1561 |
| ANO003 | 45.119 | -93.238 | 1 | 0.3117 | LAKEW001 | 48.700 | -94.561 | 2 | 0.1746 |
| BECK005 | 46.683 | -95.688 | 5 | 0.1567 | LAKEW002 | 48.713 | -94.651 | 3 | 0.0349 |
| BECK006 | 46.949 | -95.331 | 3 | 0.1283 | LAKEW003 | 48.712 | -94.596 | 3 | 0.2827 |
| BECK008 | 46.921 | -95.259 | 3 | 0.1413 | MAH001 | 47.375 | -95.603 | 3 | 0.0649 |
| BECK009 | 46.827 | -95.690 | 3 | 0.0038 | MAH002 | 47.178 | -95.714 | 2 | 0.0552 |
| BECK010 | 47.013 | -95.341 | 5 | 0.1431 | MARS001 | 48.266 | -95.933 | 2 | 0.2636 |
| BECK011 | 47.043 | -95.278 | 3 | 0.2809 | MOR003 | 46.070 | -93.893 | 3 | 0.2626 |
| BECK012 | 46.829 | -96.132 | 4 | 0.0700 | PEN001 | 47.993 | -96.289 | 2 | 0.2678 |
| BECK014 | 47.150 | -95.527 | 3 | -0.0016 | PINE005 | 46.062 | -92.366 | 1 | -0.0448 |
| BEL001 | 47.929 | -94.447 | 4 | 0.1140 | PINE006 | 46.090 | -92.467 | 2 | -0.0175 |
| BEL003 | 47.900 | -94.587 | 4 | 0.2163 | POLK001 | 47.778 | -96.631 | 2 | 0.1629 |

| | | | | | | | | | |
|---|---|---|---|---|---|---|---|---|---|
| BEL004 | 47.880 | -94.907 | 4 | 0.0329 | POLK003 | 47.982 | -96.483 | 2 | 0.2826 |
| BEL005 | 47.878 | -95.239 | 3 | 0.0953 | RAM002 | 45.074 | -93.020 | 1 | 0.1493 |
| BEL006 | 48.141 | -95.254 | 5 | 0.3115 | REDL001 | 47.845 | -96.004 | 3 | 0.3027 |
| BEL009 | 48.282 | -95.336 | 2 | 0.1252 | ROS001 | 48.953 | -95.600 | 2 | 0.2597 |
| BEL011 | 47.560 | -94.835 | 3 | 0.3744 | ROS003 | 48.716 | -94.736 | 2 | 0.1570 |
| BEL012 | 47.654 | -94.678 | 5 | 0.1260 | ROS004 | 48.713 | -95.455 | 2 | 0.1176 |
| BEL013 | 47.730 | -94.585 | 3 | 0.1132 | ROS005 | 48.663 | -95.342 | 3 | 0.3487 |
| BEL015 | 47.472 | -94.480 | 4 | 0.0808 | ROS006 | 48.698 | -94.952 | 1 | 0.1601 |
| BEL017 | 47.702 | -95.000 | 4 | 0.2763 | SHER003 | 45.411 | -93.686 | 2 | 0.2491 |
| BEL018 | 47.862 | -95.155 | 4 | 0.2247 | STEAR004 | 46.072 | -94.483 | 2 | -0.0562 |
| BEL019 | 48.000 | -95.252 | 3 | 0.3127 | STEV003 | 45.471 | -96.130 | 3 | -0.0562 |
| BEL020 | 47.850 | -94.815 | 5 | 0.3096 | STL001 | 46.715 | -92.205 | 4 | 0.0145 |
| BEL021 | 47.874 | -94.438 | 3 | 0.0386 | STL002 | 46.718 | -92.037 | 5 | 0.2277 |
| BEN001 | 45.576 | -94.126 | 3 | 0.3737 | STL004 | 46.774 | -92.141 | 4 | 0.0160 |
| CASS002 | 47.374 | -94.546 | 5 | 0.1778 | STL005 | 46.816 | -92.100 | 4 | 0.0039 |
| CASS007 | 46.716 | -94.406 | 5 | 0.1286 | STL006 | 47.740 | -91.486 | 4 | 0.0693 |
| CASS008-15 | 46.543 | -94.687 | 1 | 0.3310 | STL007 | 46.836 | -92.014 | 5 | 0.0228 |
| CASS008-16 | 47.337 | -94.623 | 3 | 0.1424 | STL008 | 46.885 | -91.912 | 3 | -0.0743 |
| CASS008-18 | 47.057 | -93.919 | 5 | 0.0842 | STL009 | 47.807 | -91.887 | 2 | 0.0518 |
| CLEAR001 | 47.525 | -95.508 | 4 | 0.0981 | STL010 | 47.816 | -92.237 | 2 | 0.0119 |
| CLEAR002 | 47.833 | -95.427 | 4 | 0.2073 | STL012 | 47.950 | -92.829 | 4 | 0.2590 |
| CLEAR003 | 47.359 | -95.383 | 3 | -0.0154 | STL013 | 46.928 | -92.918 | 4 | -0.0981 |
| CLEAR004 | 47.146 | -95.560 | 4 | 0.0708 | STL014 | 47.051 | -92.950 | 3 | 0.0664 |

| | | | | | | | | | |
|---|---|---|---|---|---|---|---|---|---|
| CLEAR005 | 47.253 | -95.223 | 5 | 0.2895 | STL015 | 47.268 | -92.023 | 4 | -0.0205 |
| COOK001 | 47.577 | -90.830 | 4 | 0.0072 | STL016 | 47.406 | -92.963 | 5 | 0.0878 |
| COOK002 | 47.830 | -89.983 | 2 | -0.1054 | STL017 | 47.435 | -92.921 | 5 | -0.0680 |
| CROW005 | 46.272 | -93.855 | 2 | 0.0979 | STL018 | 47.480 | -92.871 | 5 | 0.1223 |
| CROW006 | 46.380 | -94.246 | 4 | 0.2337 | STL019 | 47.674 | -92.866 | 4 | 0.0521 |
| CROW007 | 46.344 | -94.192 | 5 | 0.2630 | STL020 | 47.777 | -92.569 | 5 | 0.3904 |
| HUB002 | 47.217 | -94.748 | 4 | 0.0121 | STL021 | 48.050 | -92.832 | 5 | 0.1543 |
| HUB003 | 46.978 | -95.204 | 4 | 0.0697 | STL022 | 48.109 | -92.840 | 4 | 0.2873 |
| HUB004 | 46.926 | -95.052 | 5 | 0.1224 | STL024 | 47.769 | -92.654 | 4 | 0.1096 |
| HUB006 | 47.006 | -94.725 | 4 | 0.0603 | STL025 | 47.643 | -92.560 | 5 | 0.1064 |
| HUB008 | 46.760 | -94.777 | 1 | 0.1004 | STL026 | 47.593 | -92.561 | 5 | 0.0025 |
| HUB009-15 | 46.971 | -94.724 | 4 | 0.0926 | STL027 | 47.294 | -94.477 | 4 | 0.0816 |
| HUB009-16 | 47.310 | -95.067 | 3 | 0.0228 | STL028 | 47.087 | -92.474 | 4 | -0.0519 |
| HUB010-15 | 47.167 | -94.796 | 3 | 0.2975 | STL029 | 46.910 | -92.387 | 5 | 0.0105 |
| HUB010-16 | 47.483 | -95.130 | 4 | 0.0667 | STL030 | 46.972 | -92.218 | 5 | 0.1876 |
| HUB011 | 47.189 | -95.052 | 4 | 0.1018 | STL031 | 47.393 | -92.811 | 4 | 0.0128 |
| HUB012 | 47.249 | -95.191 | 5 | 0.2204 | STL032 | 47.459 | -92.389 | 4 | 0.0328 |
| HUB013 | 47.011 | -95.106 | 3 | 0.0281 | STL033 | 47.625 | -92.244 | 4 | -0.0868 |
| ISA003 | 45.572 | -93.209 | 3 | 0.2200 | STL034 | 47.800 | -92.283 | 5 | 0.0416 |
| ISA005 | 45.443 | -93.235 | 2 | 0.2399 | STL036 | 47.412 | -91.977 | 4 | 0.0971 |
| ITA002 | 47.823 | -94.265 | 5 | 0.0548 | STL038 | 47.606 | -91.646 | 3 | 0.3200 |
| ITA003 | 47.743 | -94.055 | 3 | 0.0900 | STL039 | 47.371 | -92.037 | 3 | 0.1574 |
| ITA004 | 47.727 | -94.007 | 4 | 0.0465 | STL040 | 47.372 | -92.310 | 5 | 0.1725 |

| | | | | | | | | | |
|---|---|---|---|---|---|---|---|---|---|
| ITA005 | 47.760 | -93.723 | 4 | 0.1785 | SWI004 | 45.334 | -95.963 | 1 | 0.0472 |
| ITA006 | 47.807 | -93.384 | 3 | 0.1010 | SWI005 | 45.357 | -95.337 | 1 | 0.0156 |
| ITA008 | 47.517 | -93.194 | 4 | 0.0787 | WAB004 | 44.382 | -92.038 | 1 | 0.2328 |
| ITA009 | 47.402 | -93.174 | 5 | 0.0793 | WAB005 | 44.360 | -92.367 | 1 | 0.1955 |
| ITA010 | 47.323 | -93.276 | 5 | 0.1427 | WASH004 | 45.101 | -92.985 | 1 | 0.1746 |
| ITA011 | 47.291 | -93.421 | 5 | 0.2427 | WASH005 | 45.238 | -92.990 | 2 | 0.0136 |
| ITA012 | 47.301 | -93.417 | 3 | -0.0945 | WIN017 | 43.938 | -91.861 | 1 | 0.0301 |

Experimental Design Details

**Figure S1. Experimental Plot Layout**

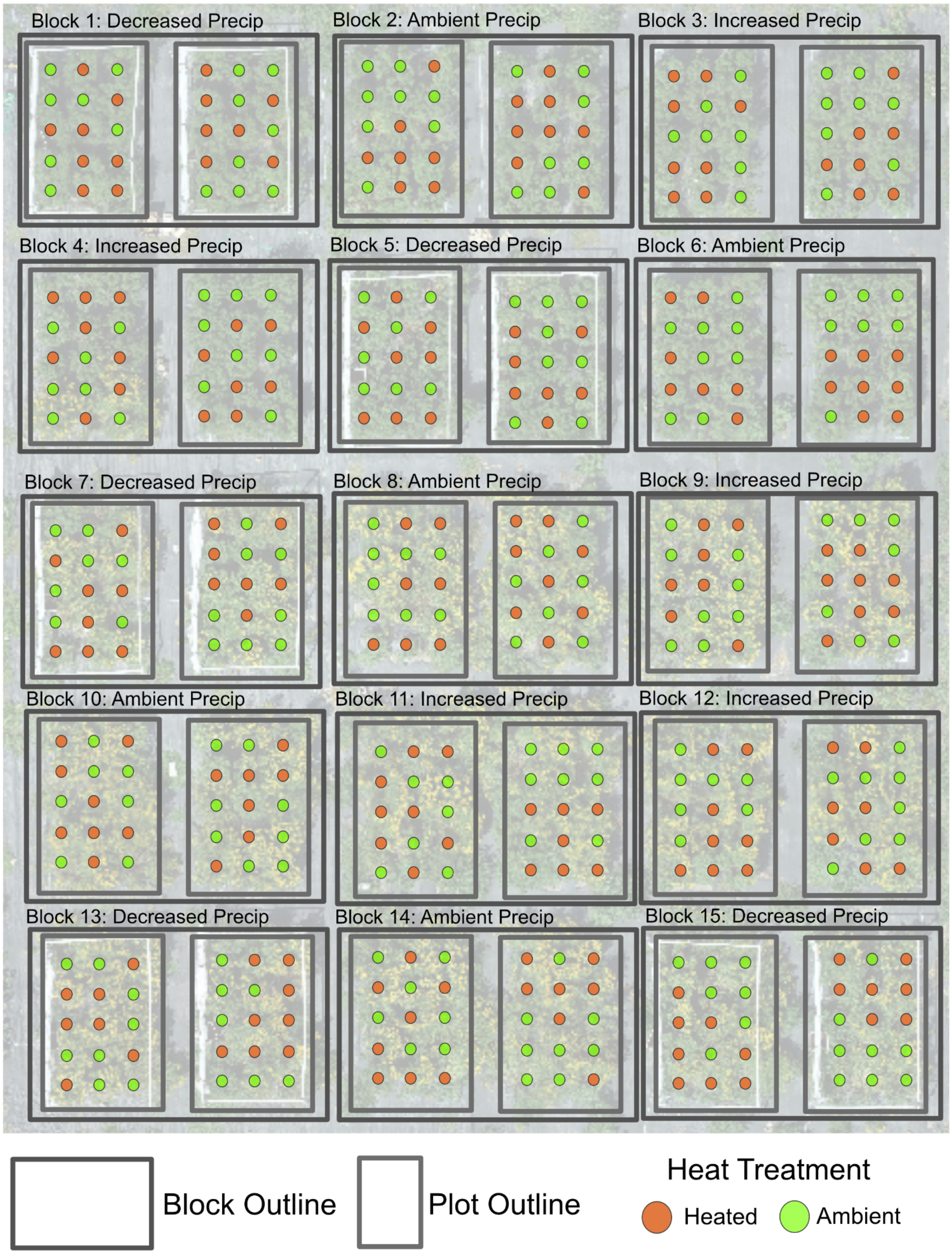

#### Planting

We started experimental plants in the greenhouse. Seeds were planted individually in 120 well seedling trays in damp promix soil and kept on a misting bench for one week. After the week, plants were moved to a greenhouse. Once plants had reached the 1-2 leaf stage, they were moved outside to benches in a partially shaded area to harden off for transplantation. Seedlings were transplanted into field plots at the 2-4 leaf stage on 30 June 2021. We watered transplants for a week to limit death from transplant shock. When seedling stocks allowed, we replaced plants that had died in the first two weeks and watered for one week post transplantation.

To prevent unintended spread of common tansy from our experimental garden, we implemented two containment strategies: we covered all unplanted areas with black weed cloth and transplanted experimental seedlings into nursery pots (8” diameter x 10” depth) that were recessed into the ground using a soil auger. We refilled pots with native field soil.

#### Climate manipulations

OTCs were constructed out of transparent corrugated greenhouse roofing materials cut into 1.5’ x 2’ panels that were zip-tied together to form a rectangular prism with an open top and bottom. Three weeks post planting (7/19/2021-7/20/2021), we secured OTCs around heat-treated plants using stakes threaded through the zip ties at the four corners of the OTCs. OTCs were maintained around treated plants for the duration of the experiment, including winter. We tracked thermal conditions using 20 iButton loggers for the first season. Data loggers were placed at soil level (n = 20 (10/ treatment level)). Loggers were protected by a waterproof plasticine coating and nested within ventilated, solar-shielded (white-painted) Falcon tubes. These assemblies were mounted on pin flags to ensure consistent positioning adjacent to the study individuals.

**Figure S2. OTC Temperature manipulations**

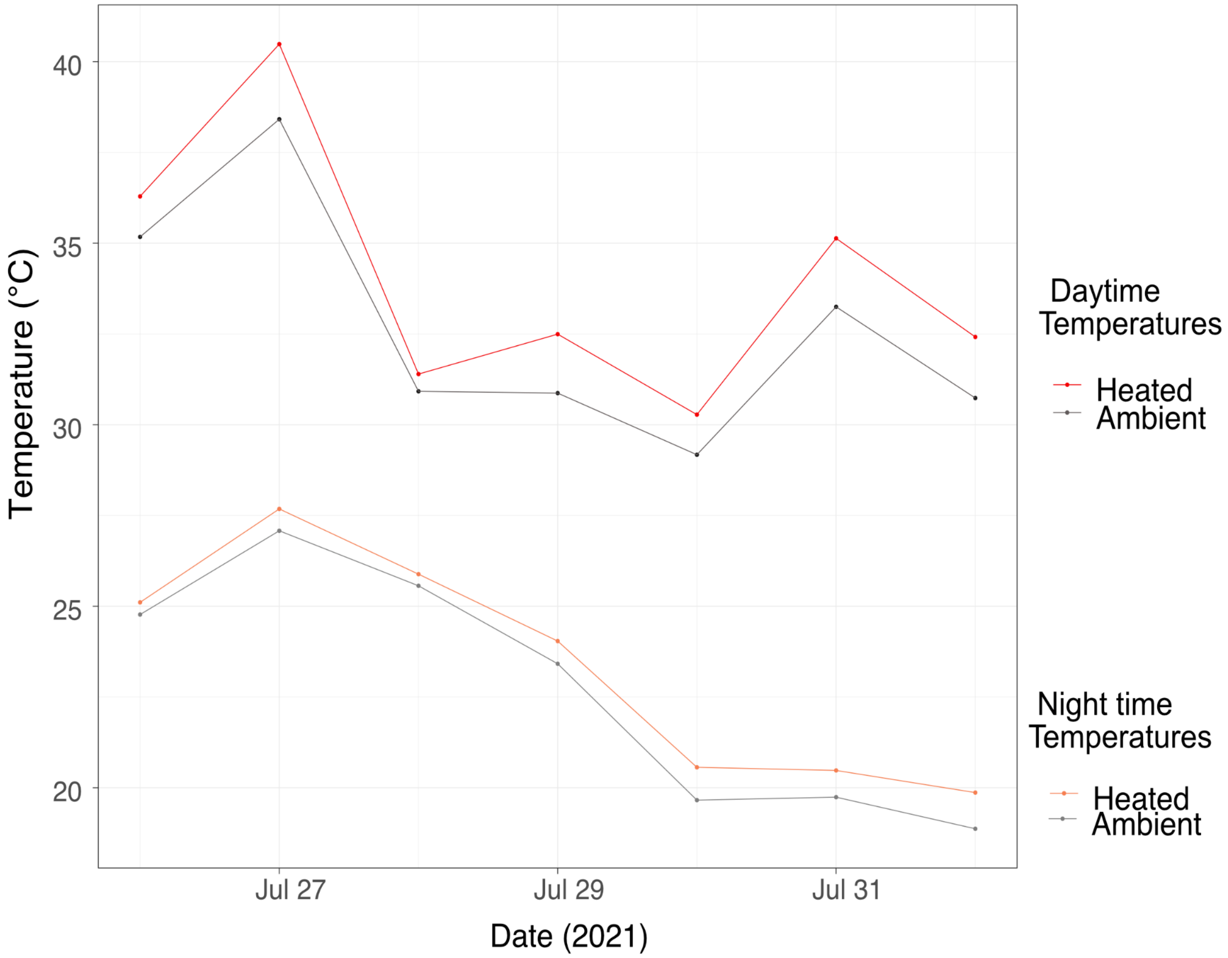


We excluded precipitation using event-based rain-out shelters. During targeted events, approximately every third rain event, we covered plots with plastic sheeting secured to 1.75 m PVC frames. This system funneled water into barrels and piped it away from the experimental area. Exclusion decisions were based on cumulative precipitation estimates from the campus weather station (https://www.dnr.state.mn.us/climate/climate_monitor/climate_observatory.html). For precipitation additions, we added water to simulate a 30% increase within 24 hours (generally <12) after each rain event.

#### Figure S3. Precipitation manipulations

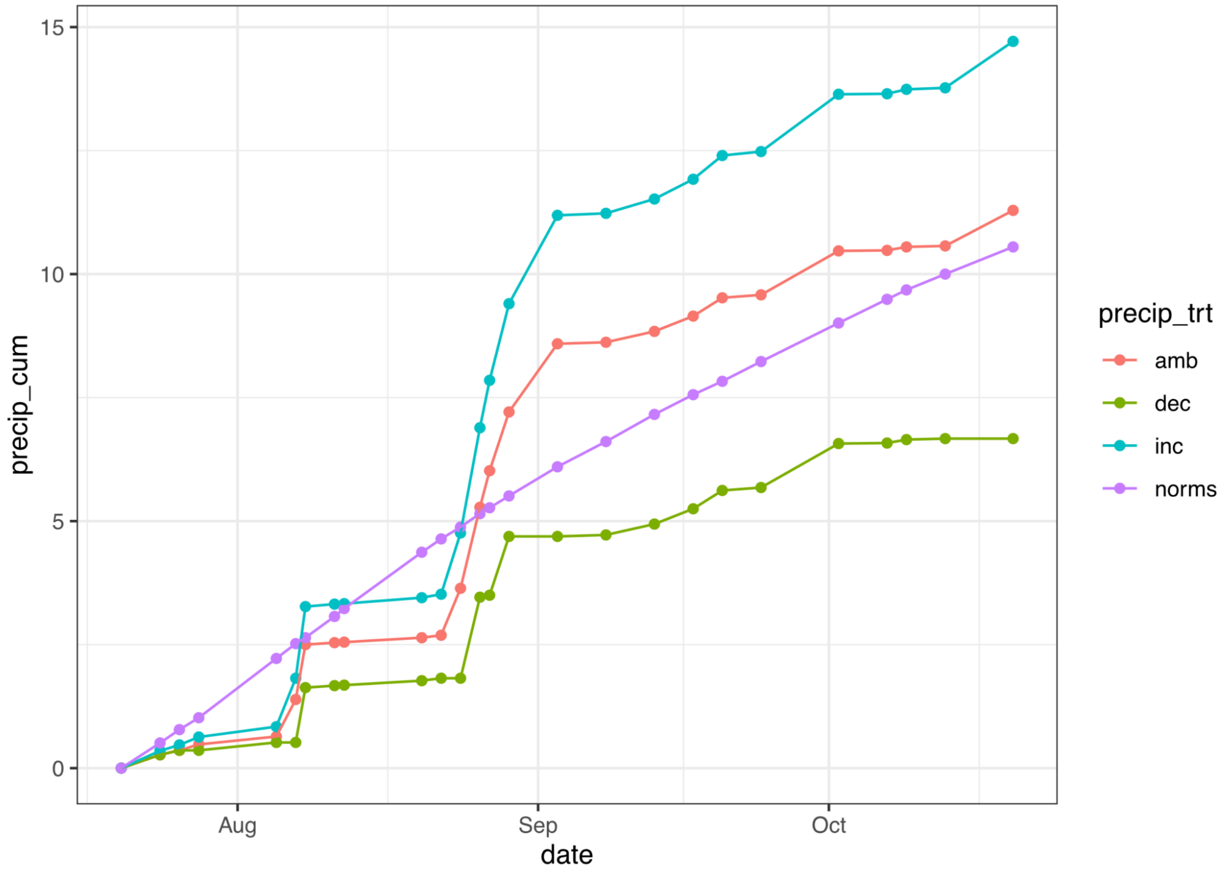


Detailed fitness PCs

#### Figure S4. Composite fitness

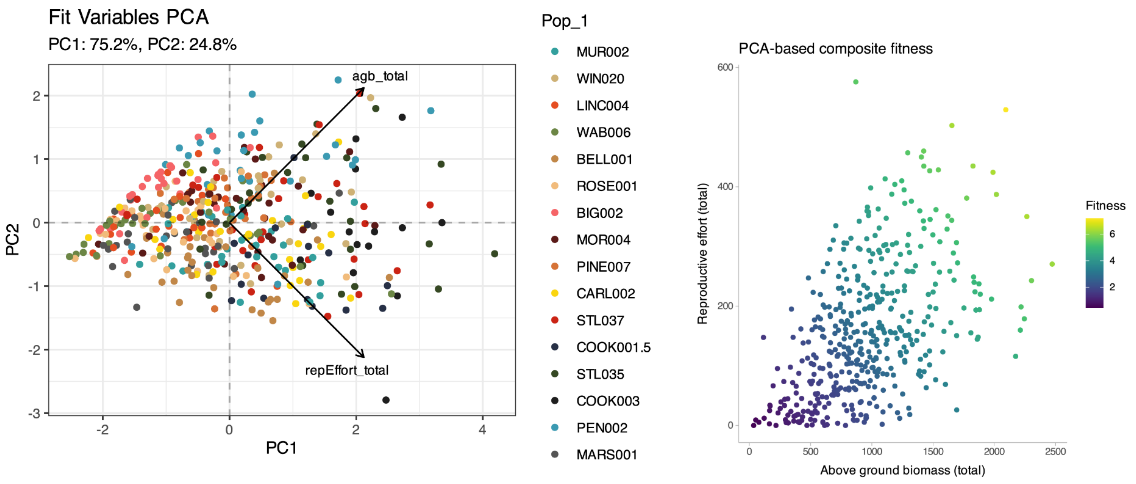

### Climate Variables

#### Table S2. Chelsea Climate Variables

| Variable | Full Name | Unit | Description | Subset |
|---|---|---|---|---|
| Bio1 | Mean Annual Temperature | °C | Mean annual temperature calculated as the average of mean monthly temperatures over the year | Temperature |
| Bio12 | Cumulative Annual Precipitation | kg m-2 year-1 | Sum of monthly precipitation totals across the year | Precipitation |
| CMI_mean | Mean Climate Moisture Index | kg m-2 | Average of monthly ratio of precipitation to potential evapotranspiration; indicator of climatic water availability | Precipitation |
| CMI_range | Range of Climate Moisture Index | kg m-2 | Annual range of average CMI values | Precipitation |
| Frost_Change_Freq | Frost Change Frequency | count | Number of freeze–thaw transitions per year | Temperature |
| First_Day_Grow_Seas | First Day of the Growing Season (TREECLIM) | julian day | Julian day marking the first occurrence of growing season conditions | Temperature |
| GDD10 | Growing Degree Days (10°C) | °C | Sum of daily mean temperatures above 10 °C accumulated over the year | Temperature |
| Grow_Seas_Length | Growing Season Length (Days) | days | Number of days between the first and last occurrence of growing season conditions | Temperature |
| Grow_Seas_Precip | Growing Season Precipitation | kg m-2 gsl-1 | Total precipitation accumulated during the growing season period | Precipitation |
| Grow_Seas_Temp | Growing Season Temperature | °C | Average daily mean temperature over all growing season days | Temperature |
| Num_GDD10 | Number of Growing Degree Days at (10°C) | number of days | Total number of days in a year with mean daily temperature above 10 °C | Temperature |
| PET_mean | Mean Potential Evapotranspiration | kg m-2 | Average of total potential evapotranspiration for the month assuming unlimited water availability calculated based on Penman Monteith | Temperature |
| PET_range | Range of Potential Evapotranspiration | | Range of the average monthly potential evapotranspiration | Temperature |

**Figure S5. Chelsa ClimateVariables**

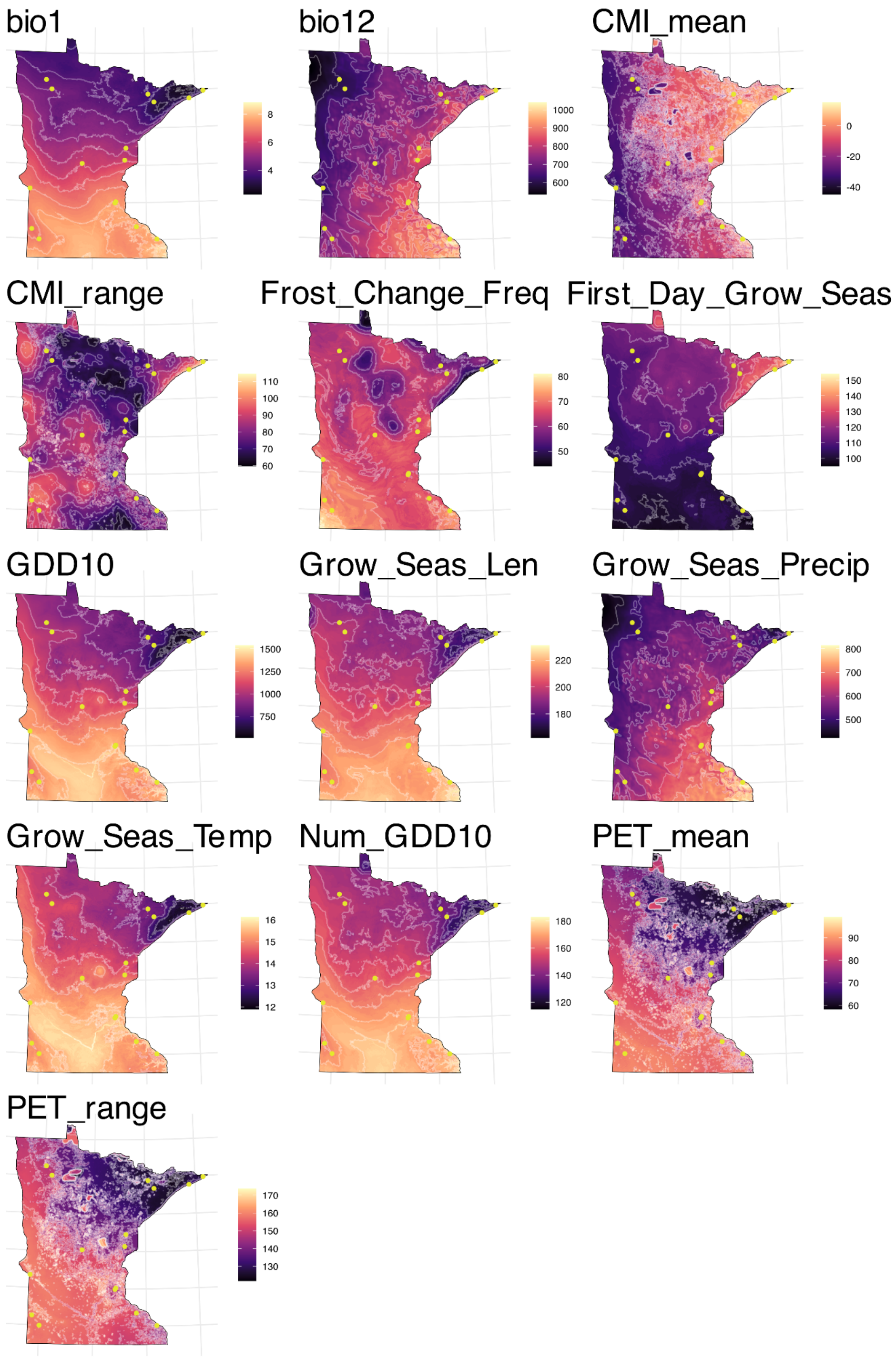

**Table S3. Distance Correlations**

| | Distance to invasion core | Environmental distance (all variables) | Environmental distance (temp. variables) |
|---|---|---|---|
| Environmental distance (all variables) | -0.495 | | |
| Environmental distance (temperature variables) | -0.695** | 0.951*** | |
| Environmental distance (precipitation variables) | 0.015 | 0.819*** | 0.607* |

#### Estimates of Population Mean Fitness

For population-level analyses and figures, we extracted population-level estimated marginal means (EMM) of fitness from M0 by averaging the model's predictions across all combinations of temperature and precipitation treatments within each population. To generate confidence intervals, we propagated the full coefficient covariance matrix using the linear predictor matrix (lpmatrix in mgcv). We accounted for the fact that smoothing parameters are themselves estimated with uncertainty by calculating the unconditional variance-covariance matrix (unconditional = TRUE) when estimating error. The random block intercept was excluded from EMM prediction so that the resulting estimates are marginal over both treatments and blocks.

### Supplementary Results

##### Figure S6. Observed population mean fitness and fitness variation

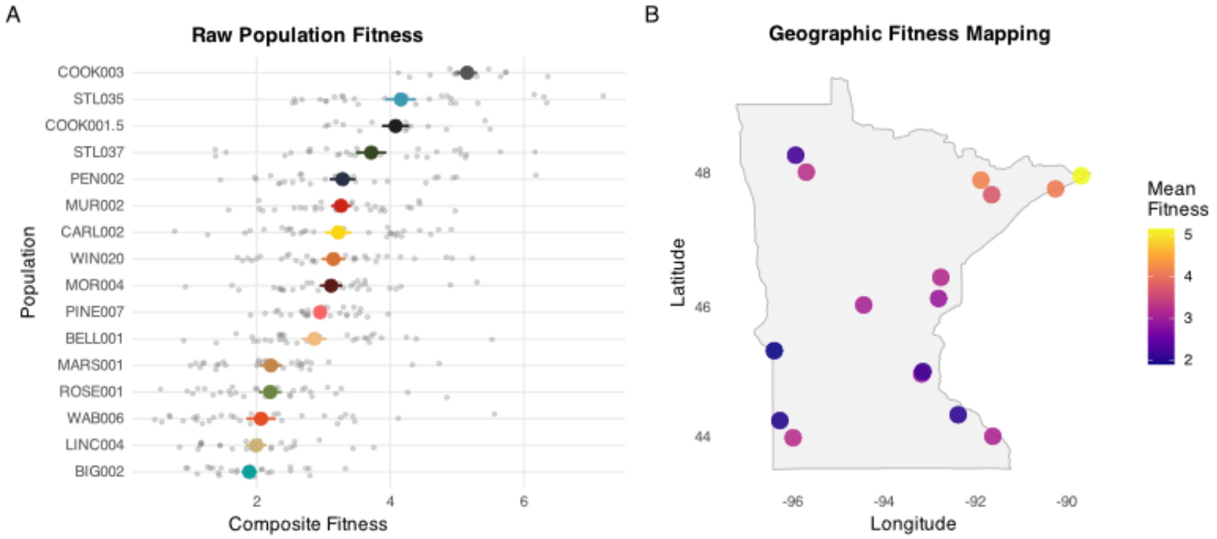

**Table S4. GAM effects table**

| Term | Est./ edf | SE/ Ref df | Stat (t/F) | P |
|---|---|---|---|---|
| **M0_null** | | | | |
| Parametric | | | | |
| (Intercept) | 3.08 | 0.22 | 13.96 | <0.001 |
| HeatTrt.L | -0.27 | 0.08 | -3.48 | <0.001 |
| PrecipTrt.L | -0.05 | 0.11 | -0.49 | 0.6243 |
| PrecipTrt.Q | 0.04 | 0.11 | 0.34 | 0.7356 |
| HeatTrt.L:PrecipTrt.L | 0.08 | 0.11 | 0.80 | 0.4258 |
| HeatTrt.L:PrecipTrt.Q | -0.12 | 0.11 | -1.12 | 0.2615 |
| Smooth | | | | |
| s(Pop_1) | 13.99 | 15 | 29.80 | <0.001 |
| s(Pop_1,HeatTrt) | 6.00 | 30 | 0.48 | 0.1686 |
| s(Pop_1,PrecipTrt) | 0.01 | 45 | 0.00 | 0.5831 |
| s(Block) | 6.43 | 12 | 1.49 | 0.0018 |
| **M1_LA_full** | | | | |
| Parametric | | | | |
| (Intercept) | 3.00 | 0.16 | 19.04 | <0.001 |
| HeatTrt.L | -0.27 | 0.08 | -3.46 | <0.001 |
| PrecipTrt.L | -0.05 | 0.11 | -0.49 | 0.6261 |
| PrecipTrt.Q | 0.04 | 0.11 | 0.33 | 0.7408 |
| HeatTrt.L:PrecipTrt.L | 0.08 | 0.11 | 0.80 | 0.4266 |
| HeatTrt.L:PrecipTrt.Q | -0.12 | 0.11 | -1.13 | 0.2583 |
| Smooth | | | | |
| s(s_env_distance) | 1.83 | 1.87 | 10.00 | 0.002 |
| s(Pop_1) | 11.33 | 14 | 14.66 | <0.001 |
| s(Pop_1,HeatTrt) | 6.35 | 29 | 0.54 | 0.061 |
| s(Pop_1,PrecipTrt) | 0.01 | 44 | 0.00 | 0.5853 |
| s(Block) | 6.40 | 12 | 1.40 | 0.0027 |
| **M2_LA_temp** | | | | |
| Parametric | | | | |
| (Intercept) | 3.01 | 0.15 | 20.39 | <0.001 |
| HeatTrt.L | -0.27 | 0.08 | -3.47 | <0.001 |
| PrecipTrt.L | -0.05 | 0.11 | -0.50 | 0.6201 |
| PrecipTrt.Q | 0.04 | 0.11 | 0.34 | 0.7369 |
| HeatTrt.L:PrecipTrt.L | 0.08 | 0.11 | 0.79 | 0.4272 |
| HeatTrt.L:PrecipTrt.Q | -0.12 | 0.11 | -1.13 | 0.2589 |
| Smooth | | | | |
| s(s_env_distance_temp) | 1.00 | 1 | 21.97 | <0.001 |
| s(Pop_1) | 11.75 | 14 | 13.85 | <0.001 |
| s(Pop_1,HeatTrt) | 6.49 | 29 | 0.56 | 0.0665 |
| s(Pop_1,PrecipTrt) | 0.00 | 44 | 0.00 | 0.5812 |

| Term | Est./ edf | SE/ Ref df | Stat (t/F) | P |
|---|---|---|---|---|
| **M4.H_EL** | | | | |
| Parametric | | | | |
| (Intercept) | 3.05 | 0.16 | 18.59 | <0.001 |
| HeatTrt.L | -0.27 | 0.08 | -3.59 | <0.001 |
| PrecipTrt.L | -0.06 | 0.11 | -0.51 | 0.6133 |
| PrecipTrt.Q | 0.04 | 0.11 | 0.35 | 0.7265 |
| HeatTrt.L:PrecipTrt.L | 0.08 | 0.11 | 0.80 | 0.4265 |
| HeatTrt.L:PrecipTrt.Q | -0.12 | 0.11 | -1.12 | 0.2642 |
| Smooth | | | | |
| s(s_dist_to_core) | 1.00 | 1 | 14.34 | <0.001 |
| s(s_dist_to_core):HeatTrtHeat | 1.00 | 1 | 0.79 | 0.3757 |
| s(Pop_1) | 12.33 | 14 | 15.69 | <0.001 |
| s(Pop_1,HeatTrt) | 5.25 | 28 | 0.43 | 0.1041 |
| s(Pop_1,PrecipTrt) | 0.00 | 44 | 0.00 | 0.5799 |
| s(Block) | 6.41 | 12 | 1.41 | 0.0025 |
| **M4.P_EL** | | | | |
| Parametric | | | | |
| (Intercept) | 3.05 | 0.16 | 18.51 | <0.001 |
| HeatTrt.L | -0.27 | 0.08 | -3.47 | <0.001 |
| PrecipTrt.L | -0.05 | 0.11 | -0.49 | 0.6227 |
| PrecipTrt.Q | 0.04 | 0.11 | 0.35 | 0.7293 |
| HeatTrt.L:PrecipTrt.L | 0.08 | 0.11 | 0.80 | 0.426 |
| HeatTrt.L:PrecipTrt.Q | -0.12 | 0.11 | -1.12 | 0.2653 |
| Smooth | | | | |
| s(s_dist_to_core) | 1.00 | 1 | 9.46 | 0.0022 |
| s(s_dist_to_core):PrecipTrtDec | 1.00 | 1 | 0.20 | 0.6526 |
| s(s_dist_to_core):PrecipTrtInc | 1.00 | 1 | 1.34 | 0.2486 |
| s(Pop_1) | 12.24 | 14 | 16.44 | <0.001 |
| s(Pop_1,HeatTrt) | 6.29 | 29 | 0.52 | 0.1137 |
| s(Pop_1,PrecipTrt) | 0.00 | 42 | 0.00 | 0.546 |
| s(Block) | 6.46 | 12 | 1.43 | 0.0025 |
| **M5_joint_full** | | | | |
| Parametric | | | | |
| (Intercept) | 3.02 | 0.15 | 20.51 | <0.001 |
| HeatTrt.L | -0.27 | 0.08 | -3.48 | <0.001 |
| PrecipTrt.L | -0.06 | 0.11 | -0.50 | 0.6147 |
| PrecipTrt.Q | 0.04 | 0.11 | 0.35 | 0.7275 |
| HeatTrt.L:PrecipTrt.L | 0.08 | 0.11 | 0.80 | 0.4256 |
| HeatTrt.L:PrecipTrt.Q | -0.12 | 0.11 | -1.12 | 0.2634 |
| Smooth | | | | |

| | | | | |
|---|---|---|---|---|
| s(Block) | 6.43 | 12 | 1.41 | 0.0026 |
| **M2.H_LA_temp_xHeat** | | | | |
| Parametric | | | | |
| (Intercept) | 3.01 | 0.15 | 20.38 | <0.001 |
| HeatTrt.L | -0.27 | 0.08 | -3.55 | <0.001 |
| PrecipTrt.L | -0.06 | 0.11 | -0.50 | 0.6193 |
| PrecipTrt.Q | 0.04 | 0.11 | 0.34 | 0.7348 |
| HeatTrt.L:PrecipTrt.L | 0.08 | 0.11 | 0.79 | 0.4321 |
| HeatTrt.L:PrecipTrt.Q | -0.12 | 0.11 | -1.12 | 0.2626 |
| Smooth | | | | |
| s(s_env_distance_temp) | 1.00 | 1 | 21.32 | <0.001 |
| s(s_env_distance_temp):HeatTrtHeat | 1.00 | 1 | 0.53 | 0.4683 |
| s(Pop_1) | 11.87 | 14 | 13.36 | <0.001 |
| s(Pop_1,HeatTrt) | 5.53 | 28 | 0.47 | 0.068 |
| s(Pop_1,PrecipTrt) | 0.00 | 44 | 0.00 | 0.5845 |
| s(Block) | 6.43 | 12 | 1.41 | 0.0026 |
| **M3_LA_precip** | | | | |
| Parametric | | | | |
| (Intercept) | 3.05 | 0.21 | 14.29 | <0.001 |
| HeatTrt.L | -0.27 | 0.08 | -3.47 | <0.001 |
| PrecipTrt.L | -0.05 | 0.11 | -0.49 | 0.624 |
| PrecipTrt.Q | 0.04 | 0.11 | 0.34 | 0.7363 |
| HeatTrt.L:PrecipTrt.L | 0.08 | 0.11 | 0.80 | 0.4261 |
| HeatTrt.L:PrecipTrt.Q | -0.12 | 0.11 | -1.13 | 0.2609 |
| Smooth | | | | |
| s(s_env_distance_prec) | 1.00 | 1 | 1.35 | 0.2464 |
| s(Pop_1) | 12.98 | 14 | 31.09 | <0.001 |
| s(Pop_1,HeatTrt) | 6.02 | 29 | 0.51 | 0.1197 |
| s(Pop_1,PrecipTrt) | 0.01 | 44 | 0.00 | 0.5826 |
| s(Block) | 6.44 | 12 | 1.48 | 0.0019 |
| **M3.P_LA_precip_xPrecip** | | | | |
| Parametric | | | | |
| (Intercept) | 3.05 | 0.21 | 14.28 | <0.001 |
| HeatTrt.L | -0.27 | 0.08 | -3.46 | <0.001 |
| PrecipTrt.L | -0.05 | 0.11 | -0.49 | 0.6233 |
| PrecipTrt.Q | 0.04 | 0.11 | 0.34 | 0.7348 |
| HeatTrt.L:PrecipTrt.L | 0.08 | 0.11 | 0.79 | 0.43 |
| HeatTrt.L:PrecipTrt.Q | -0.12 | 0.11 | -1.13 | 0.2605 |
| Smooth | | | | |
| s(s_env_distance_prec) | 1.00 | 1 | 1.93 | 0.1659 |
| s(s_env_distance_prec):PrecipTrtDec | 1.00 | 1 | 0.09 | 0.7584 |
| s(s_env_distance_prec):PrecipTrtInc | 1.00 | 1 | 1.78 | 0.1832 |
| s(Pop_1) | 12.97 | 14 | 31.07 | <0.001 |
| s(Pop_1,HeatTrt) | 6.06 | 29 | 0.52 | 0.1216 |
| s(Pop_1,PrecipTrt) | 0.00 | 42 | 0.00 | 0.6192 |
| s(Block) | 6.50 | 12 | 1.50 | 0.0018 |
| **M4_EL** | | | | |
| Parametric | | | | |
| (Intercept) | 3.05 | 0.16 | 18.60 | <0.001 |
| HeatTrt.L | -0.27 | 0.08 | -3.48 | <0.001 |
| PrecipTrt.L | -0.06 | 0.11 | -0.50 | 0.6154 |
| PrecipTrt.Q | 0.04 | 0.11 | 0.35 | 0.7249 |

| | | | | |
|---|---|---|---|---|
| s(s_env_distance) | 1.00 | 1 | 4.61 | 0.0324 |
| s(s_dist_to_core) | 1.00 | 1 | 6.46 | 0.0114 |
| s(Pop_1) | 10.90 | 13 | 14.30 | <0.001 |
| s(Pop_1,HeatTrt) | 6.42 | 28 | 0.57 | 0.0598 |
| s(Pop_1,PrecipTrt) | 0.01 | 43 | 0.00 | 0.5736 |
| s(Block) | 6.43 | 12 | 1.41 | 0.0026 |
| **M6_joint_temp** | | | | |
| Parametric | | | | |
| (Intercept) | 3.01 | 0.14 | 21.46 | <0.001 |
| HeatTrt.L | -0.27 | 0.08 | -3.47 | <0.001 |
| PrecipTrt.L | -0.06 | 0.11 | -0.50 | 0.6158 |
| PrecipTrt.Q | 0.04 | 0.11 | 0.34 | 0.7304 |
| HeatTrt.L:PrecipTrt.L | 0.08 | 0.11 | 0.80 | 0.4262 |
| HeatTrt.L:PrecipTrt.Q | -0.12 | 0.11 | -1.12 | 0.2617 |
| Smooth | | | | |
| s(s_env_distance_temp) | 1.00 | 1 | 6.90 | 0.0089 |
| s(s_dist_to_core) | 1.00 | 1 | 1.80 | 0.1806 |
| s(Pop_1) | 10.67 | 13 | 12.76 | <0.001 |
| s(Pop_1,HeatTrt) | 6.51 | 28 | 0.58 | 0.0535 |
| s(Pop_1,PrecipTrt) | 0.00 | 43 | 0.00 | 0.5731 |
| s(Block) | 6.43 | 12 | 1.40 | 0.0026 |
| **M6.H_joint_temp_xHeat** | | | | |
| Parametric | | | | |
| (Intercept) | 3.01 | 0.14 | 21.45 | <0.001 |
| HeatTrt.L | -0.27 | 0.08 | -3.55 | <0.001 |
| PrecipTrt.L | -0.06 | 0.11 | -0.50 | 0.615 |
| PrecipTrt.Q | 0.04 | 0.11 | 0.35 | 0.7284 |
| HeatTrt.L:PrecipTrt.L | 0.08 | 0.11 | 0.79 | 0.4312 |
| HeatTrt.L:PrecipTrt.Q | -0.12 | 0.11 | -1.12 | 0.2655 |
| Smooth | | | | |
| s(s_env_distance_temp) | 1.00 | 1 | 7.42 | 0.0067 |
| s(s_env_distance_temp):HeatTrtHeat | 1.00 | 1 | 0.54 | 0.4649 |
| s(s_dist_to_core) | 1.00 | 1 | 1.80 | 0.1799 |
| s(Pop_1) | 10.79 | 13 | 12.34 | <0.001 |
| s(Pop_1,HeatTrt) | 5.53 | 27 | 0.49 | 0.0563 |
| s(Pop_1,PrecipTrt) | 0.00 | 43 | 0.00 | 0.5764 |
| s(Block) | 6.42 | 12 | 1.40 | 0.0027 |
| **M7_joint_precip** | | | | |
| Parametric | | | | |
| (Intercept) | 3.03 | 0.15 | 19.67 | <0.001 |
| HeatTrt.L | -0.27 | 0.08 | -3.48 | <0.001 |
| PrecipTrt.L | -0.06 | 0.11 | -0.51 | 0.6135 |
| PrecipTrt.Q | 0.04 | 0.11 | 0.35 | 0.7248 |
| HeatTrt.L:PrecipTrt.L | 0.08 | 0.11 | 0.80 | 0.425 |
| HeatTrt.L:PrecipTrt.Q | -0.12 | 0.11 | -1.12 | 0.2648 |
| Smooth | | | | |
| s(s_env_distance_prec) | 1.00 | 1 | 2.71 | 0.1005 |
| s(s_dist_to_core) | 1.00 | 1 | 16.02 | <0.001 |
| s(Pop_1) | 11.09 | 13 | 15.81 | <0.001 |
| s(Pop_1,HeatTrt) | 6.34 | 28 | 0.56 | 0.0667 |
| s(Pop_1,PrecipTrt) | 0.01 | 43 | 0.00 | 0.5739 |
| s(Block) | 6.43 | 12 | 1.42 | 0.0025 |

| | | | | |
|---|---|---|---|---|
| HeatTrt.L:PrecipTrt.L | 0.08 | 0.11 | 0.80 | 0.4245 |
| HeatTrt.L:PrecipTrt.Q | -0.12 | 0.11 | -1.12 | 0.2654 |
| Smooth | | | | |
| s(s_dist_to_core) | 1.00 | 1 | 13.69 | <0.001 |
| s(Pop_1) | 12.23 | 14 | 16.37 | <0.001 |
| s(Pop_1,HeatTrt) | 6.27 | 29 | 0.52 | 0.1103 |
| s(Pop_1,PrecipTrt) | 0.01 | 44 | 0.00 | 0.5759 |
| s(Block) | 6.42 | 12 | 1.42 | 0.0025 |

| **M7.P_joint_precip_xPrecip** | | | | |
|---|---|---|---|---|
| Parametric | | | | |
| (Intercept) | 3.03 | 0.15 | 19.69 | <0.001 |
| HeatTrt.L | -0.27 | 0.08 | -3.47 | <0.001 |
| PrecipTrt.L | -0.06 | 0.11 | -0.51 | 0.613 |
| PrecipTrt.Q | 0.04 | 0.11 | 0.35 | 0.7233 |
| HeatTrt.L:PrecipTrt.L | 0.08 | 0.11 | 0.79 | 0.429 |
| HeatTrt.L:PrecipTrt.Q | -0.12 | 0.11 | -1.12 | 0.2647 |
| Smooth | | | | |
| s(s_env_distance_prec) | 1.00 | 1 | 3.65 | 0.0568 |
| s(s_env_distance_prec):PrecipTrtDec | 1.00 | 1 | 0.13 | 0.723 |
| s(s_env_distance_prec):PrecipTrtInc | 1.00 | 1 | 1.87 | 0.1717 |
| s(s_dist_to_core) | 1.00 | 1 | 16.08 | <0.001 |
| s(Pop_1) | 11.07 | 13 | 15.72 | <0.001 |
| s(Pop_1,HeatTrt) | 6.39 | 28 | 0.57 | 0.0675 |
| s(Pop_1,PrecipTrt) | 0.00 | 41 | 0.00 | 0.6122 |
| s(Block) | 6.50 | 12 | 1.44 | 0.0023 |

**Figure S7. Best joint GAM relationships**

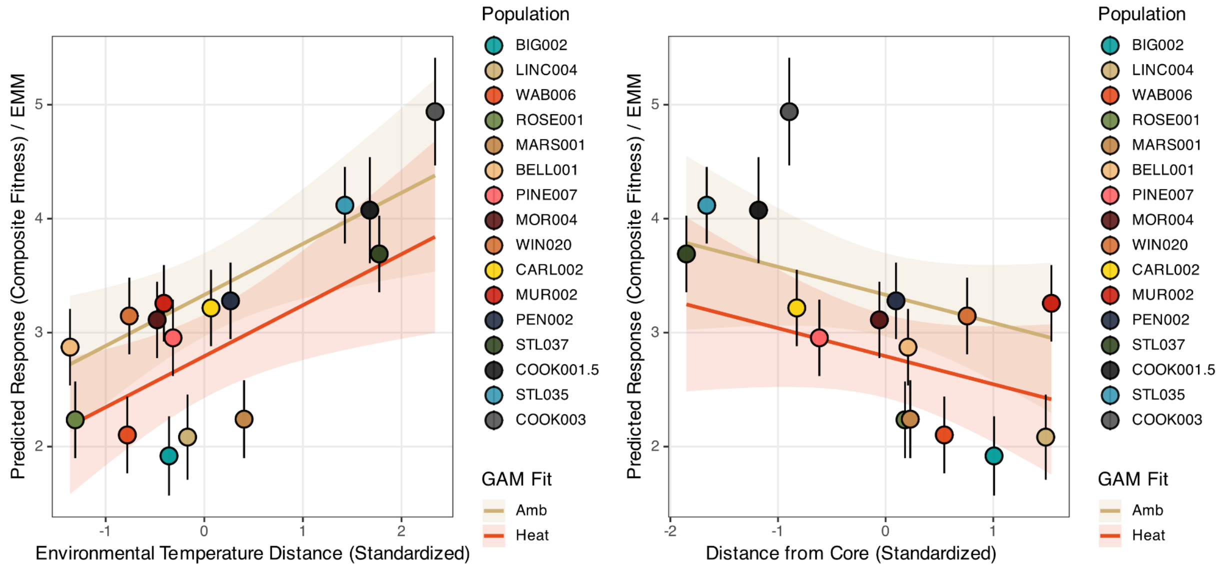


**Table S5. Correlation among expansion load factors excluding outliers**

| | | | |
|---|---|---|---|
| MN Common Tansy Populations | | | |
| | Distance to core | Population Size Category | F |
| Population Size Category | **-0.511** | | |
| F | **0.267** | -0.028 | 1 |
| Experimental Populations | | | |
| | Distance to core | Population Size Category | F |
| Population Size Category | **-0.732** | | |
| F | **0.805** | -0.513 | |
| Estimated Mean Fitness | **-0.662** | **0.843** | -0.505 |

**Table S6. Correlations among expansion load factors including outliers**

| | Distance to core | Population Size Category |
|---|---|---|
| Population Size Category | **-0.519** | |
| F | **0.183** | -0.052 |

**Figure S8. Components of expansion load with outliers**

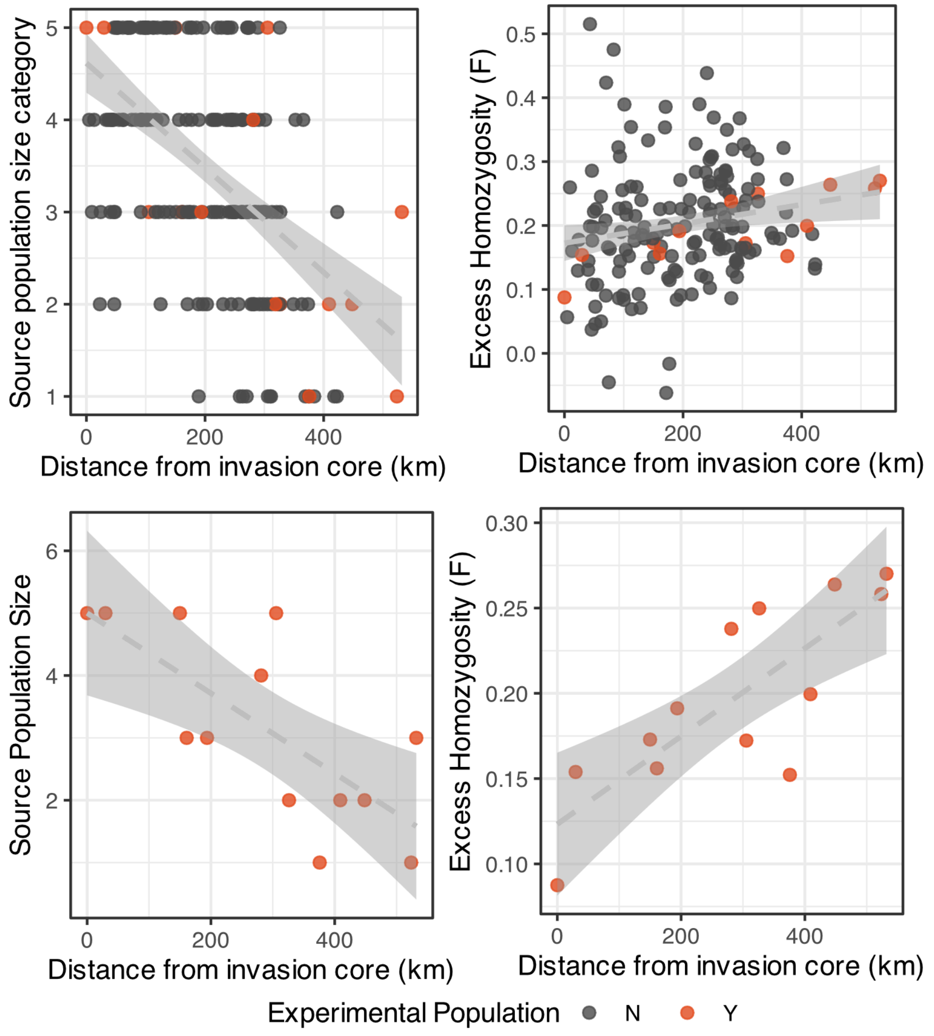
